# Contrasting functions of *Arabidopsis* SUMO1/2 isoforms with SUMO3 intersect to modulate innate immunity and global SUMOylome responses

**DOI:** 10.1101/2020.05.15.097535

**Authors:** Kishor Dnyaneshwar Ingole, Mritunjay Kasera, Harrold A. van den Burg, Saikat Bhattacharjee

## Abstract

Reversible covalent attachment of SMALL UBIQUITIN-LIKE MODIFIERS (SUMOs) on target proteins regulate diverse cellular process across all eukaryotes. In *Arabidopsis thaliana*, most mutants with perturbed global SUMOylome display severe impairments in growth and adaptations to physiological stresses. Since SUMOs self-regulate activities of SUMOylation-associated proteins, existence of multiple isoforms introduces possibilities of their functional intersections which remain unexplored especially in plant systems. Using well-established defense responses elicited against virulent and avirulent *Pseudomonas syringae* pv. *tomato* strains, we investigated crosstalks in individual and combinatorial Arabidopsis *sum* mutants. Here we report that while *SUM1* and *SUM2* additively, but not equivalently suppress basal and TNL-specific immunity via down-regulation of salicylic acid (SA)-dependent responses, *SUM3* promotes these defenses genetically downstream of SA. Remarkably, the expression of *SUM3* is transcriptionally suppressed by SUMO1 or SUMO2. The loss of *SUM3* not only lowers basal or post-bacterial challenge responsive enhancements of SUMO1/2-congugates but also reduces upregulation dynamics of defensive proteins and SUMOylation-associated transcripts. Combining a *sum3* mutation partially attenuates heightened immunity of *sum1* or *sum2* mutants suggesting intricate functional impingements among these isoforms in optimizing immune amplitudes. Similar *SUM1*-*SUM3* intersections also affect global SUMOylome responses to heat-shock affecting most notably the induction of selective heat-shock transcription factors. Overall, our investigations reveal novel insights into auto-regulatory mechanisms among SUMO isoforms in host SUMOylome maintenance and adjustments to environmental challenges.

**Author Summary:** In plants, similar to animals, protein functions are regulated at multiple levels. One prevalent mode is to allow covalent linkage of small proteins to specific amino acids on targets thereby affecting its fate and function. One such kind of modification named as SUMOylation involves attachment of SUMO proteins. A plant maintains strict control over its pool of SUMOylated proteins (termed SUMOylome) which upon biotic or abiotic stresses are altered to facilitate appropriate responses, returning back to steady-state when the threat subsides. In mutants of the model plant *Arabidopsis thaliana* having disturbed steady-state SUMOylome, growth and developmental defects ensue. These mutants are auto-immune showing more resistance to infection by the bacterial pathogen *Pseudomonas syringae*. However, Arabidopsis SUMO-family are comprised of multiple members raising the question about their specificity or functional crosstalks. We discovered that two SUMO members function in coordination to suppress immunity including the repression of a third member which supports defenses. The expression of this third member during pathogen attack or heat-shock influences the responsive changes in the host SUMOylome likely suggesting SUMOs themselves play vital role in these adaptations. Overall, our work highlights novel intersections of SUMO members in mounting stress-specific responses.

## Introduction

Post-translational modifications (PTMs) dynamically and reversibly modulate target proteins properties either by affecting their fate, location, or function and facilitate transitory adaptations often necessary for developmental passages and responding to biotic/abiotic cues. Of special importance in this category is the reversible attachment of the SMALL UBIQUITIN-LIKE MODIFIER (SUMO) through a process known as SUMOylation. This highly conserved mode of covalently modifying proteins is utilized primarily to regulate nuclear functions such as DNA replication, chromatin remodeling, transcription, and post-transcriptional processes in eukaryotes [1–3]. In plants, temperature perturbations, salinity changes and application of defensive hormones such as salicylic acid (SA) cause massive changes on global SUMOylome [4–6]. Interestingly, a vast majority of differentially SUMOylated proteins are nuclear, perhaps indicative of a rapid mechanism to modulate transcriptome in response to stimulus. Computational studies identify SUMOylation-annotated proteins as central relay players of protein-protein interaction webs, especially for transcriptional processes [7].

SUMOylation cascade recruits processed SUMOs with C-terminus di-glycine (GG) residues, a product of catalysis by specialized SUMO or Ubiquitin-like proteases (ULPs), to covalently attach via an isopeptide bond to ε-amino group of lysine residues on target proteins. The modified lysine often is a part of a partially conserved motif, Ψ-K-X-D/E (Ψ = hydrophobic amino acid, X= any amino acid, D/E= aspartate/glutamate). A conjugatable SUMO, subsequently forms a thiol-ester bond with the SUMO E1 ACTIVATING ENZYME (SAE). A trans-esterification reaction further shuttles SUMO to the SUMO E2 CONJUGATING ENZYME 1 (SCE1) and then to target lysine on SUMOylation substrates. Enrichment of SUMOylated proteins upon heat shock reveal that SUMO1 can conjugate to itself and form poly-SUMO1 chains [5]. Although SCE1 is capable of catalyzing polySUMO-chain formation, members of SUMO E4 ligases the PROTEIN INHIBITOR OF ACTIVATED STAT LIKE protein (PIAL1/2 in Arabidopsis) enhance polySUMOylation [2,8,9]. Substrate specificities both for mono- or polySUMOylation are further regulated by SUMO E3 LIGASES such as HIGH PLOIDY2/METHYL METHANE SULFONATE21 (HPY2/MMS21) and SAP and MIZ1 (SIZ1) [10]. Fate of covalently-attached SUMOs may either include de-conjugation and recycling by SUMO proteases or targeted proteolysis of the substrate through ubiquitin-mediated pathway. Indeed, proteasome components are enriched in poly-SUMO1 pull-downs [5]. These interactions are non-covalent in nature facilitated by hydrophobic amino acid-rich SUMO-interaction motifs (SIMs) present in the recipient. A group of moderately conserved SUMO-targeted ubiquitin ligases (STUbLs) bind poly-SUMO chains via internal SIMs to ubiquitinylate the substrate [11]. Interestingly, SIMs are also only enriched in the SUMOylation-associated proteins such as SIZ1, SCE1, SAE2, SUMO-protease EARLY IN SHORT DAYS 4 (ESD4), PIAL1/2, STUbLs implying strong auto-regulatory mechanisms [8,12]. Not the least, increasing evidences that reciprocal influences of phosphorylation, ubiquitination, and acetylation that often compete with or modulate the efficacy of SUMOylation highlights the complexity of crosstalks among PTMs [13–15].

Unlike fruit fly, worm, or yeast, humans and plants express multiple SUMO isoforms [16]. In *Arabidopsis thaliana*, 4 SUMO isoforms are expressed namely SUMO1, SUMO2, SUMO3 and SUMO5 [4,17]. SUMO1 and SUMO2 share 89% sequence identity, whereas SUMO3 and SUMO5 are considerably diverged from SUMO1 (48% and 35% identical, respectively). The homolog pair SUMO1/2 are a result of genomic duplication event of a SUMO clade that preceded the evolution of monocots and eudicots from common ancient angiosperms [18]. Tandem organization of *SUM2* and *SUM3* genes in Arabidopsis are a result of gene duplication subsequently followed by diversification of only *SUM3* sequences. At relative expression levels, *SUM1/2* are more abundant than *SUM3* or *SUM5* [19]. Partial overlaps in tissue-specific expression patterns and biochemical properties taken together with embryonic lethality of *sum1 sum2* double mutant indicate that plants require at least one functional copy of either of these redundant isoforms [20–22]. The *Arabidopsis sum3* mutant is viable with mild late-flowering phenotype [21]. Acutely different SUMO3 however, unlike SUMO1/2, cannot form poly-SUMO chains *in vitro*, and shows little or no change to heat-shock treatment [4,17,19,21,23]. Not the least, SUMO1/2 but not SUMO3-modified targets, are efficiently deconjugated by SUMO proteases *in vitro* [17,24].

Host SUMOylome readjustments play a vital role in regulation of plant immunity and been comprehensively highlighted in several excellent reviews [25–27]. This was first evident in the Arabidopsis SUMO E3 ligase *SIZ1* mutant (*siz1-2*) and subsequently in SUMO protease mutants of *OVERLY TOLERANT TO SALT 1/2 (OTS1/2)* and *ESD4* [6,20,28,29]. Increased accumulation of defensive hormone salicylic acid (SA) and constitutive expression of PATHOGENESIS-RELATED (PR) proteins in these mutants conferred enhanced resistance when challenged with the bacterial pathogen *Pseudomonas syringae* pv. *tomato* strain DC3000 (*PstDC3000*). In this context, it is therefore not surprising that pathogens attempt to manipulate host SUMOylome to increase their colonization efficiencies [30–32]. Several bacterial phytopathogenic effectors interfere with host SUMOylation as a mode to suppress immunity [33,34]. XopD, a secreted effector from *Xanthomonas campestris pv. vesicatoria* (*Xcv*) de-conjugates SUMO from unknown targets in plants [35]. Mutations that disrupt SUMO-protease functions not only render the cognate strain deficient in virulence but also lower host defense induction.

Suggestively, both SUM1/2 jointly suppress SA-dependent defenses since plants null for *SUM1* and expressing microRNA-silenced *SUM2* (*sum1-1 amiR-SUM2*) have drastically reduced total SUMO1/2-conjugates compared to wild-type and display heightened anti-bacterial immunity [21]. The SUMO E3 ligase mutant *siz1-2*, also with lower levels of SUMO1/2 conjugates are similarly enhanced resistant to *PstDC3000* [20,28]. On the contrary, *esd4-1* or *ots1ots2* plants with deficient SUMO protease ESD4 or OTS1/2 functions, respectively have increased SUMO1/2-conjugates yet display elevated SA-dependent defenses [1,6,29]. Constitutive activation of SA-dependent defenses also occur in *SUM1/2*-overexpressing transgenic plants regardless of whether the over-expressed SUMO isoforms are conjugation-proficient or deficient [21]. Thus, it is likely that perturbations rather than increase or decrease in SUMO1/2-conjugates *per se* regulate immune responses. Unlike *SUM1/2*, *SUM3* is SA-inducible, upregulated in *sum1-1 amiR-SUM2* plants, and upon over-expression enhance the resistance to *PstDC3000* [21]. SUMO3-mediated SUMOylation of NPR1 (NONEXPRESSOR OF PATHOGENESIS-RELATED 1), a master transducer of SA-signaling is essential for defenses thus placing *SUM3* as a *bona fide* positive immune regulator [36]. Whether increased SUMO3 activities during defenses intersect to alter SUMOylome outcomes of a host remains unexplored. Role of *SUM3* in other stress responses similarly is also unknown. Evidently, perturbing host SUMOylome during biotic/abiotic stresses or through loss/gain of a specific isoform function initiates complex signaling more especially because the SUMOs themselves moderate SUMOylation proficiencies of targets. SAE2, SCE1, SIZ1, and EDS4 were identified as candidates whose SUMOylation footprints change upon stress exposure [5]. Animal studies also elegantly demonstrate ‘SUMO-preference/switching’ wherein the isoform choice for substrate conjugation is modulated not only by their relative levels but also by its influence on SUMOylation machinery [37–39]. Hence, it is beyond doubt that SUMO isoforms functionally intersect when the host SUMOylome-equilibrium is disturbed. However, such evidences from plant systems are completely lacking.

In this study, we utilized defense responses to various strains of *PstDC3000* as a measure to test the contribution of individual and combinatorial *Arabidopsis* SUMO isoforms in immunity. Our data suggest that *SUM1/2* function additively, but not equally, as negative regulators of SA-driven basal and TNL (Toll-Interleukin-1 receptor-like domain)-type immunity. In contrast, *SUM3* has a more positive immune role potentiating auto-immunity that occur due to loss of *SUM1* or *SUM2*. We demonstrate that accumulation kinetics of not only defense-associated markers but also of SUMOylation-associated genes are regulated by SUMO1-SUMO3 crosstalks. We further report that global change in a host SUMOylome in response to immune activation or heat-stress is also influenced by intersections of SUMO isoform activities. Overall, our results open newer avenues to unravel role of SUMO isoforms functions in maintenance and alterations of a host SUMOylome.

## Results

### *SUM1/2* genetically function as negative immune regulators whereas *SUM3* promotes defense responses

To study individual *Arabidopsis* SUMO isoform influences on defenses, we obtained previously characterized knockout mutant lines of *sum1-1*, *sum2-1* and *sum3-1* [20,21]. Wild-type (Col-0) and mutant plants were propagated either in short day (SD) or long day conditions (LD) as indicated for respective assays. Interestingly, in all propagation regimes and especially in SD conditions, we noticed clear growth defects in *sum1-1*, but not *sum2-1* or *sum3-1* plants. The *sum1-1* mutant was developmentally dwarf with elongated leaves, reduced fresh tissue mass and increased trichomes density although with normal architecture compared to Col-0, *sum2-1* or *sum3-1* plants (Figs 1A,B; S1 Fig). Although identical *sum1-1* mutant has been utilized previously the morphological defects we observed have not been reported earlier. The reason for this discrepancy is not clear and we speculate differences in soil compositions or growth variations as possible contributing factors. Nevertheless, the phenotypic attributes of *sum1-1* were novel and we present evidences in subsequent sections that genetically link these defects solely to loss of *SUM1*.

**Fig 1:**
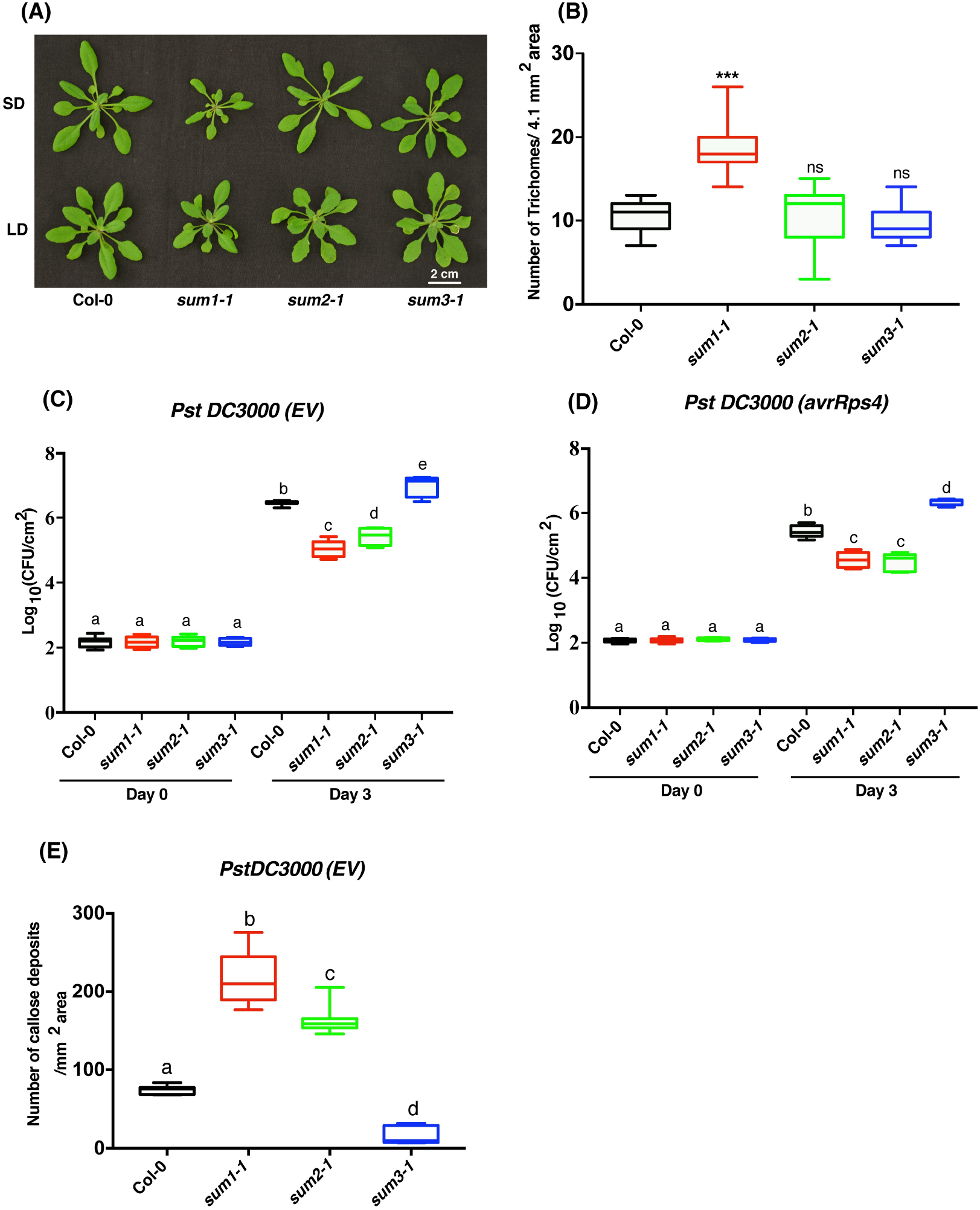
*sum1* and *sum2* mutants display enhanced whereas *sum3* is deficient in defenses against *Pseudomonas syringae* pv. *tomato* (*PstDC3000*) strains. (A) Developmental phenotypes of 4-week-old Col-0, *sum1-1, sum2-1,* and *sum3-1* mutants propagated under short day (SD) and long day (LD) growth conditions. (B) Whisker Box plot of trichome densities (#Trichomes/4.1 mm^2^) in Col-0 and *sum1-1, sum2-1,* and *sum3-1* mutants grown under SD conditions. Trichomes in 4-week-old plants were counted after imaging leaves under bright field in a fluorescence microscope. Error bars indicate standard deviation of trichome densities in 10 random images of different leaf sections (n=10). *** indicates *p* <0.001, ns= non-significant. (C-D) Growth of (C) virulent *PstDC3000(EV)* or (D) avirulent *PstDC3000*(*avrRps4*) strains, respectively in Col-0 and *sum1-1, sum2-1,* and *sum3-1* mutants. Three to four expanded leaves from 4-week-old plants of each genotype were infiltrated with the indicated bacterial suspension at a density of 5 × 10^4^ cfu ml^−1^. Leaf discs (of predefined diameter) were punched from the infiltrated area, macerated in 10mM MgCl_2_, serially diluted and plated on appropriate antibiotic plates. Bacterial titer was calculated at 0- and 3-dpi (days post-infiltration) for the indicated bacterial infection. Whisker Box plot with Tukey test; n= 12; ANOVA was performed to measure statistical significant differences (at *p-value* <0.001) in growth of bacteria in Log_10_ scale. (E) Whisker Box plot of callose deposits/mm^2^ area of infected leaves. Infiltrated leaves at 24-hpi (hours post-infiltration) were bleached in acetic acid: ethanol solution, stained with Aniline blue and were observed under DAPI filter in a fluorescence microscope. The images were analyzed in Imaris 8.0 software. The data is representative of callose deposits median from independent random sections area (n=10). Statistical significance was determined by Student’s t-test (*** indicates at *p* <0.001.

We then investigated defense responses in the *sum* mutants utilizing the standard bacterial growth assays with virulent and avirulent *PstDC3000* strains. In Col-0, *PstDC3000* is virulent and triggers basal or PAMP-triggered immunity (PTI). In contrast, *PstDC3000* harboring *avrRps4, avrRpm1, or hopA1* effector is avirulent, triggering effector-triggered immunity (ETI), mediated by the cognate resistance gene *RPS4*, *RPM1* or *RPS6*, respectively [40]. The virulent *PstDC3000* strain we used harbors the plasmid backbone into which the avirulent effectors were cloned and hence named as *PstDC3000* (*empty vector; EV*) in subsequent sections. Fully expanded leaves of 3-4 weeks-old SD-grown plants were infiltrated with the indicated *PstDC3000* strains and the growth of the bacteria measured at 0- and 3-days post-infiltration (dpi). When compared to Col-0, total bacterial growth was reduced almost 10-fold in the *sum1-1* and ~8-fold in *sum2-1* plants whereas *sum3-1* allowed more bacterial colonization (~8-fold higher) (Fig 1C). These results implied that *SUM1/2* and *SUM3* function as negative and positive regulators of PTI, respectively. When challenged with avirulent *PstDC3000* expressing *avrRps4* or *hopA1*, two TNL-specific ETI-eliciting effectors, lower bacterial accumulation than Col-0 persisted in the *sum1-1* and *sum2-1* plants (Fig 1D; S2A Fig). The *sum3-1* plants were hyper-susceptible to these avirulent infections. Curiously, modest but consistent difference with less bacterial accumulation in *sum1-1* than *sum2-1* was observed to virulent or *PstDC3000(hopA1)* but not to *PstDC3000(avrRps4)* challenges. Further on, increased callose deposits, a well-established defensive phenomenon [41], were more pronounced in *sum1-1* (~3-fold) and *sum2* (~2-fold) than Col-0 in response to both virulent as well as avirulent (*avrRps4*) *PstDC3000* (Fig 1E; S3 Fig). Although previous studies suggest redundant roles of *SUM1/2* in defenses [21], as our results implicate that their degree of contribution however may slightly vary. Likely indicative of enhanced susceptibility to *PstDC3000*, lower levels of callose than Col-0 accumulated in *sum3-1*. Remarkably, ETI in all plants to avirulent *PstDC3000* (*avrRpm1)* remained comparable suggesting that CNL-type responses are not affected by the loss of individual SUMO isoforms (S2B Fig). Indeed, as reported earlier *avrRpm1*-mediated HR is not affected in any *sum* mutants [21]. Overall, we identify a partial redundancy between *SUM1/2* with a concomitant antagonism to *SUM3* functions in immune responses to *PstDC3000* strains.

### Basal levels of SA and SA-responsive defense markers are upregulated in *sum1-1* and *sum2-1*

Significant increases in salicylic acid, both free (SA) and glucose-conjugated (SAG), mediate signal responses against *PstDC3000* [42]. A SUMOylation-deficient and enhanced resistant *siz1-2* plants have elevated SA/SAG due to upregulated expression of SA-biosynthesis gene *SID2/ICS1* (*SALICYLIC ACID INDUCTION DEFICIENT2*/ *ISOCHORISMATE SYNTHASE1*) [21,43]. To determine whether SA/SAG perturbations reflect the immune outcomes, we measured their relative levels in the different *sum* mutants. Remarkably, both *sum1-1* and *sum2-1* plants contained significantly elevated whereas the hyper-susceptible *sum3-1* had lower SA/SAG, respectively than Col-0 (Figs 2A,B). We noted that SA/SAG levels in *sum1-1* plants were slightly higher than *sum2-1*, perhaps indicative of its modestly higher degree of basal and TNL-type immunity. Transcripts of *SID2/ICS1*, defense-associated markers *PR1* and *PR2,* and PR2 protein levels were upregulated in both *sum1-1* and *sum2-1* whereas in *sum3-1* plants these were significantly lower than Col-0 (Figs 2C,D). Likewise, increased expressions of PTI markers *FLG22-INDUCED RECEPTOR LIKE KINASE 1* (*FRK1*) and *WRKY TRANSCRIPTION FACTOR 29* (*WRKY29*) [44] than Col-0 were detected in both *sum1-1* and *sum2-1* plants (Fig 2E). Relative to Col-0, *FRK1* expression remained unaltered in *sum3-1*, whereas *WRKY29* was drastically reduced suggesting that *SUM3* promotes expression of only a subset of PTI-markers. Accumulation of a well-known SA-responsive TNL-type R protein *SUPPRESSOR OF npr1-1 CONSTITUTIVE 1* (*SNC1*) is enhanced in several auto-immune mutants [45,46]. Remarkably, both *sum1-1* and *sum2-1* displayed upregulated *SNC1* expressions whereas in *sum3-1* the transcript levels remained comparable to Col-0 (Fig 2E). Since *FRK1*, *WRKY29* and *SNC1* are SA-inducible, upregulated SA-signaling sectors likely contribute to the enhanced basal and TNL-type immunity in *sum1-1* and *sum2-1* while these defenses are deficient in *sum3-1*. Also taking into account that *SUM3* but not *SUM1* or *SUM2,* is SA-inducible [21], we reasoned that enhanced resistance in *sum1-1* or *sum2-1* may be due to elevated *SUM3* expression and its role as a positive immune regulator. Indeed, we detected ~3-and 2-fold higher *SUM3* transcripts than Col-0 in *sum1-1* and *sum2-1*, respectively (Fig 2F). Relative transcript levels of *SUM1* in *sum2-1*, or *SUM2* in *sum1-1* however remain unaltered. Overall our results identify redundant but unequal roles of *SUM1* and *SUM2* in suppressing SA-dependent defenses that include expression of PTI markers, *SNC1* and *SUM3*.

**Fig 2:**
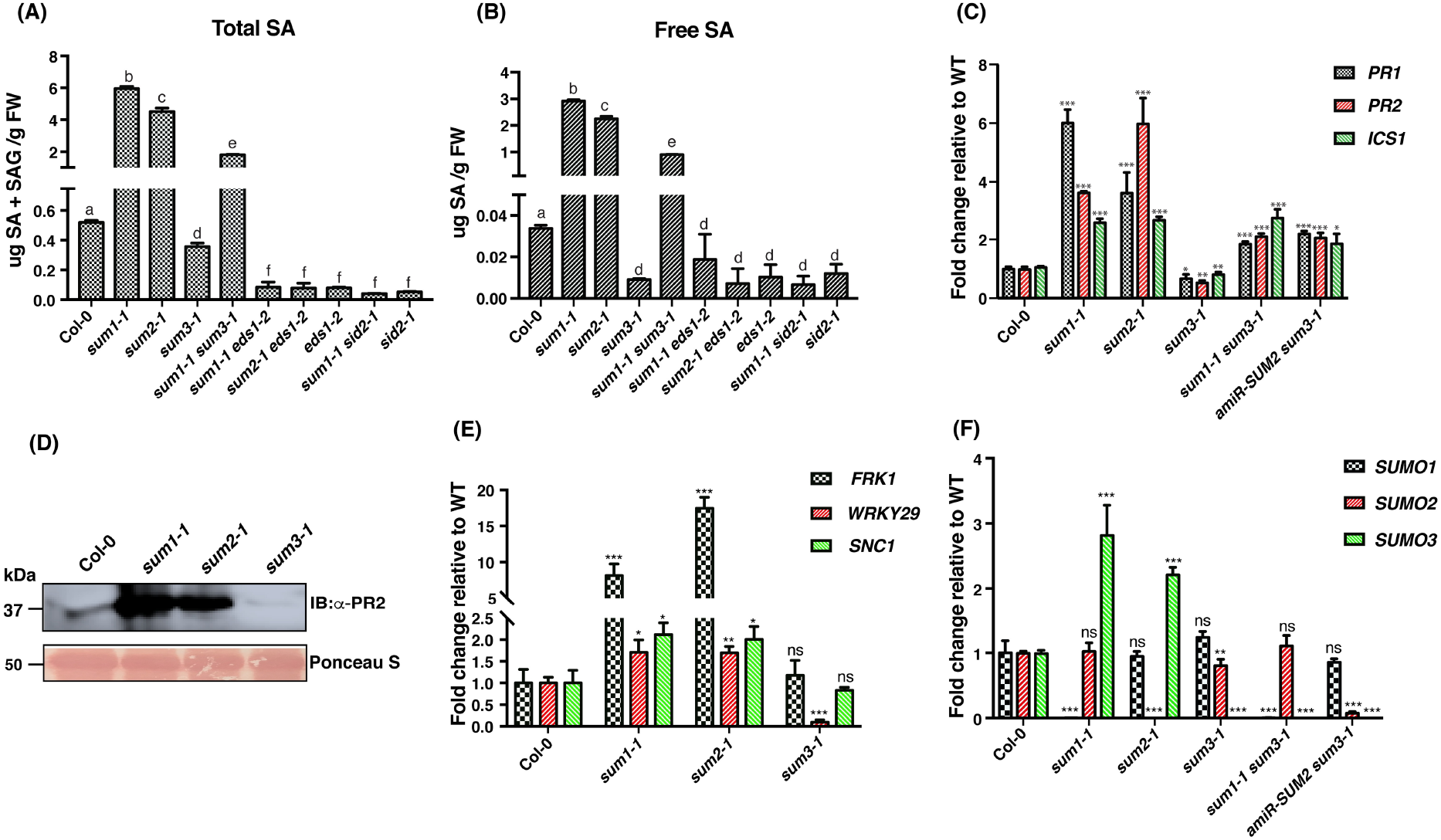
Basal SA levels and expression of several defense-associated markers are elevated in *sum1-1* or *sum2-1* whereas downregulated in *sum3-1* plants. (A) Total SA (SA + SAG) and (B) free SA levels in *sum* and combinatorial mutants in 4-week-old plants measured by biosensor *Acinetobacter* method. Data is representative of three biological replicates. Statistical significance calculated by Student’s t-test is denoted by alphabet on top of each bar. Different alphabets indicate significance at *p*<0.001. Relative transcript abundance of (C) *PR1*, *PR2*, *ICS1* (E) *FRK1*, *WRKY29*, *SNC1* (F) *SUM1*, *SUM2*, and *SUM3* in 4-week-old SD grown Col-0, *sum* and indicated combinatorial mutants was determined by qRT-PCR and normalized to *SAND* expression. The values are represented as fold change relative to Col-0 (WT). Data is representative of mean of three biological replicates (n=3). Error bars indicate SD. Student’s t-test was performed to calculate statistical significance; *=*p*<0.05; **=*p*<0.01; ***=*p*<0.001, ns= not significant. (D) Total protein extracts from 4-week-old SD grown Col-0, *sum1-1*, *sum2-1*, and *sum3-1* plants were immunoblotted with anti-PR2 antibodies. Ponceau S stain of the Rubisco subunit indicative of equal protein loading between samples is shown. Migration positions of molecular weight standards (in kDa) are indicated.

### Expression of genomic copies of *SUM1* or *SUM3* complement immune response alterations in the respective *sum* mutants

Phenotypic abnormalities of *sum1-1* plants necessitated us to genetically link the defects to loss of *SUM1*. Although our *sum1-1* mutant is identical to earlier reports [20,21], to validate our observations, we utilized the transgenic *His-H89R-SUM1* line that express a His_6_-tagged genomic *SUM1* variant (H89R) from its native promoter in a *sum1-1 sum2-1* background [5]. These plants as reported earlier not only complement the lethality of double homozygous *sum1-1 sum2-1* mutant but also functionally mirror wild-type SUMO1 SUMOylome changes upon heat stress. From the *His-H89R-SUM1* plants, the *sum2-1* mutation was segregated out by generating F2 populations upon crossing with *sum1-1*. Henceforth, we termed these plants as *sum1-1:His-H89R-SUM1*. We constantly observed that growth defects we noted earlier were always linked to homozygous *sum1-1* genotypes in absence of the *His-H89R-SUM1* transgene. Thus, a functional *SUM1* abolishes the growth defects apparent for *sum1-1* plants (S4 Fig). We noted that the *sum1-1:His-H89R-SUM1* plants had slightly elevated *SUM1* (but not *SUM2*) transcripts compared to Col-0 (Fig 3A). Implicatively, as reported earlier for *SUM1* over-expression [21], SA/SAG levels although considerably reduced in comparison to the *sum1-1* parent, were still maintained at slightly higher levels than Col-0 (Fig 3B). Increased SA/SAG levels in *sum1-1:His-H89R-SUM1* plants also resulted in upregulated *SUM3* transcripts and higher PR1 proteins in comparison to Col-0 (Figs 3A,C). In pathogen growth assays using the virulent *PstDC3000(EV)*, enhanced defenses noticeable for *sum1-1* was abrogated to Col-0 levels in the *sum1-1:His-H89R-SUM1* plants (Fig 3D). Interestingly, the avirulent *PstDC3000(hopA1)* challenge although diminished enhanced resistance relative to the *sum1-1* parent, nevertheless these plants displayed stronger defenses than Col-0 (Fig 3D). This likely is attributed to higher than Col-0 levels of SA-regulated defense networks. These results unambiguously support the loss of *SUM1* as the cause of enhanced resistance and phenotypic defects in *sum1-1*.

**Fig 3.**
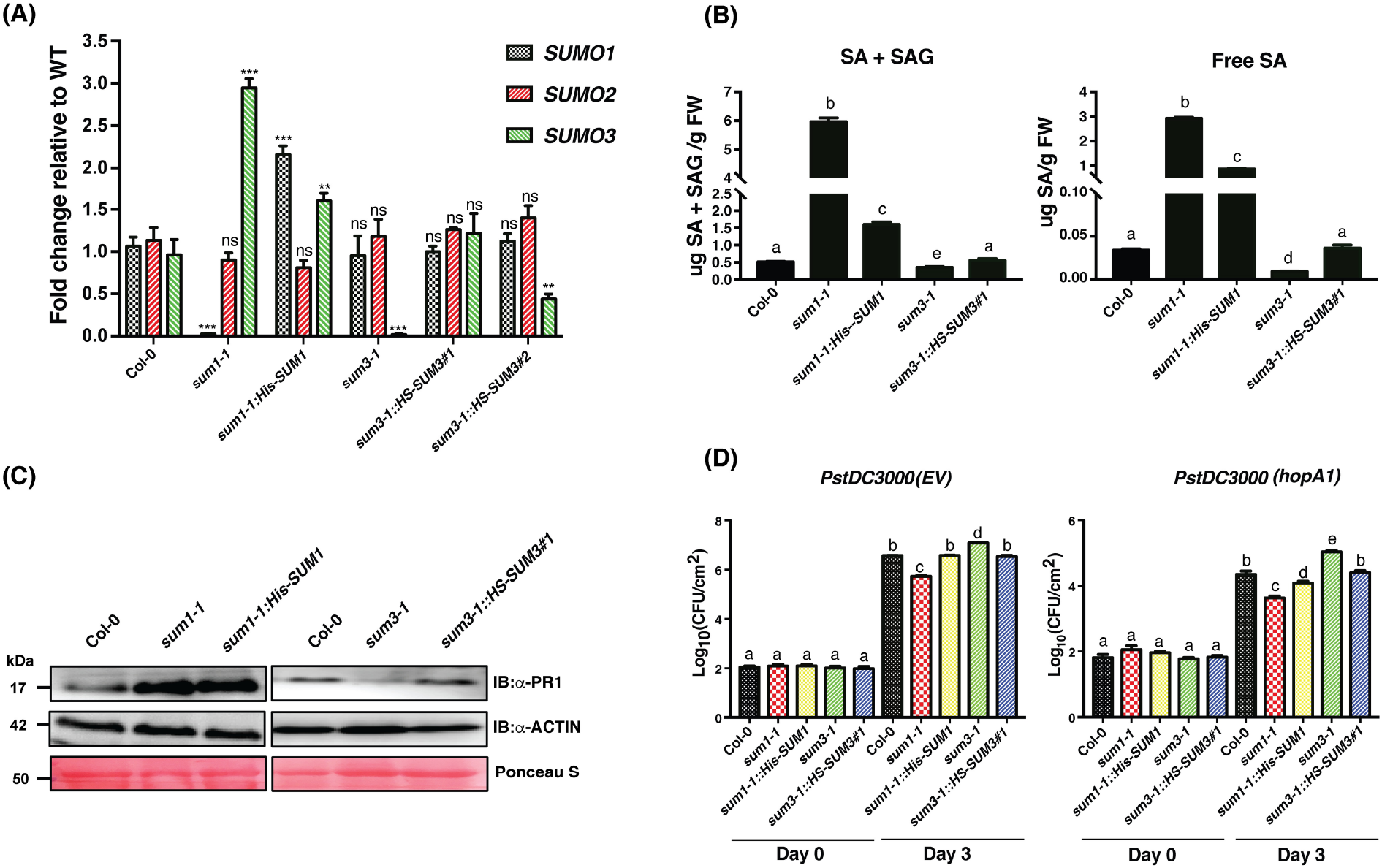
Altered defense responses to *PstDC3000* strains in *sum1-1* or *sum3-1* plants are restored to wild-type levels in the respective complemented lines. (A) Relative transcript abundance of *SUM1, SUM2* and *SUM3* in 4-week-old SD grown Col-0, *sum* mutants and the corresponding complemented line(s) was determined by qRT-PCR and normalized to *SAND* expression. The values are represented as fold change relative to Col-0 (WT). Data is representative of mean of three biological replicates (n=3). Error bars indicate SD. Statistical significance was determined by Student’s t-test; **=*p*<0.01; ***=*p*<0.001, ns= not significant. (B) Total (*left panel*) and free SA (*right panel*) levels in *sum* mutants and the complemented lines. Data is representative of three biological replicates. Statistical significance was determined by Student’s t-test and represented by alphabets with significance at *p*<0.001. (C) Endogenous PR1 protein levels in *sum* and complemented lines. Immunoblot with anti-Actin antibodies and Ponceau S staining of membrane show equal protein loading among the extracts. (D) Growth of (*left panel*) virulent *PstDC3000(EV)* or (*right panel*) avirulent *PstDC3000(hopA1)* strains, respectively in the indicated plants. Leaves from 4-week-old plants of each line were infiltrated with the indicated bacterial suspension at a density of 5 × 10^4^ cfu ml^−1^. Leaf discs from the infiltrated area was macerated in 10mM MgCl_2_, serially diluted and plated on appropriate antibiotic plates. Bacterial titers calculated at 0- and 3-dpi (days post-infiltration) for the indicated infection are shown. Different alphabets denote statistical significance at *p-value* <0.001.

We also generated two independent complemented lines of *sum3-1* to validate immune deficiencies due to the loss of *SUM3*. These transgenic plants express native promoter-driven His_6_-StrepII-tagged genomic fragment of *SUM3* (*sum3-1:HS-SUM3*). Although owing to its low abundance [21] the expression of His_6_-StrepII-SUMO3 proteins remained undetectable in immunoblots, transcripts of *SUM3* drastically reduced in the *sum3-1*, were restored to equivalent or slightly lower (~0.5X) than Col-0 levels in Line #1 and Line #2, respectively (Fig 3A). Relative abundance of *SUM1* or *SUM2* remained unaffected in both lines. For further assays, we therefore continued with *sum3-1:HS-SUM3*#1. We observed that deficiencies in SA/SAG or PR1 protein accumulations inherent to *sum3-1* were reinstated to Col-0 levels in the *sum3-1:HS-SUM3#1* plants suggesting functional complementation by the transgene (Figs 3B,C). Pathogen growth assays with either virulent *PstDC3000(EV)* or avirulent *PstDC3000(hopA1*) strains regained Col-0 levels of resistance in *sum3-1:HS-SUM3#1* (Fig 3D). These data lead us to conclude that defensive deficiencies in *sum3-1* is indeed due to loss of *SUM3*.

### Constitutively active defenses in *sum1-1* and *sum2-1* caused rapid induction of immunity

To further establish that upregulated or dampened defense marker expressions in *sum1-1*, *sum2-1* or *sum3-1*, respectively contribute to corresponding immune outcomes we challenged these plants with virulent *PstDC3000(EV)* or avirulent *PstDC3000(avrRps4*) strains. Leaf extracts harvested at 6-, 12-, and 24-hpi were used to compare accumulation kinetics of *PR1/PR2* transcripts and proteins (Fig4; S5 Fig). Basal level (0-hpi) of PR1 in Col-0, *sum2-1*, or *sum3-1* was below detection limit whereas in *sum1-1* some enhancements were noticeable (Fig 4A). At 6- or 12-hpi although Col-0 or *sum3-1* barely accumulated sufficient PR1/PR2 in response to both pathogen challenges, both *sum1-1* and *sum1-2* had markedly elevated levels of these proteins (Figs 4B,C). A similar trend continued at 24-hpi wherein in Col-0, PR2 but not PR1 proteins levels, almost matched *sum1-1* or *sum2-1*. Most significantly, even at the 24-hpi *sum3-1* plants were deficient in accumulating wild-type levels of PR1 or PR2. Real-time accumulation kinetics of *PR1*/*PR2* transcripts to both bacterial infiltrations in general mirrored the corresponding protein levels although in some instances a direct correlation was not apparent (S5 Fig). Although the reason for this discrepancy is not clear, suggested role of SUMOylation in selective translation and/or its crosstalk with other PTMs that modulate protein synthesis or stability may likely be the cause [47,48]. Nevertheless, these data provide strong molecular evidence of constitutive SA-regulated defenses in *sum1-1* and *sum2-1* conferring their enhanced resistance to *PstDC3000* strains. Contrastingly, delayed induction of defenses upon a pathogen challenge likely makes *sum3-1* hypersusceptible.

**Fig 4:**
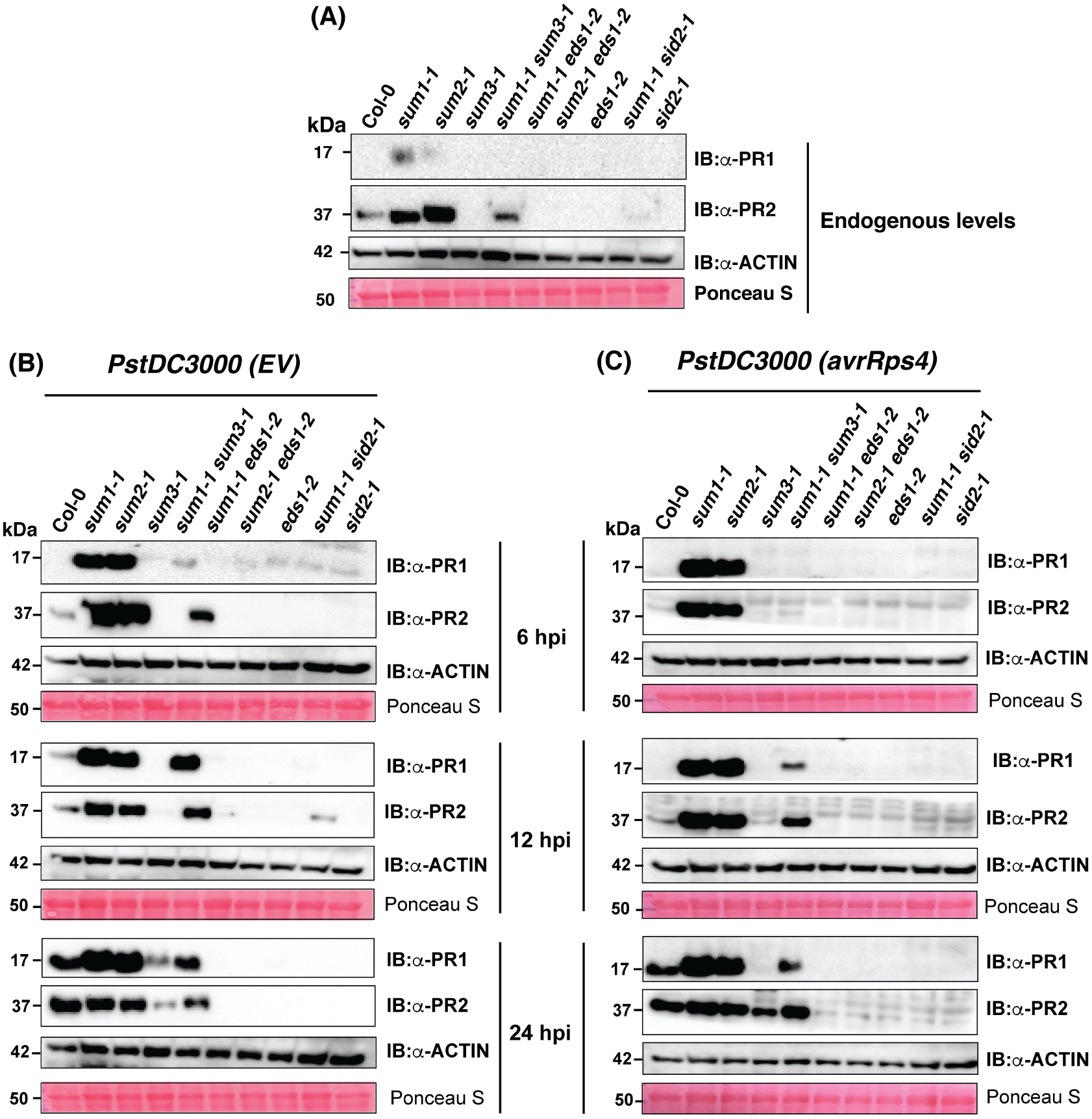
*sum1-1* and *sum2-1* display enhanced basal and rapid induction of PR1 and PR2 proteins upon pathogen challenge whereas *sum3-1* plants are deficient in these responses. Total protein from 4-week-old SD grown Col-0, *sum1-1*, *sum2-1*, *sum3-1*, *sum1-1 sum3-1*, *sum1-1 eds1-2*, *sum2-1 eds1-2*, *eds1-2*, *sum1-1 sid2-1*, and *sid2-1* was extracted from (A) un-infiltrated or at 6-, 12-, and 24-hpi (hours post-infiltration) with either (B) virulent *PstDC3000(EV)* or (C) avirulent *PstDC3000(avrRps4)* strains and immunoblotted with anti-PR1 or anti-PR2 antibodies. The membranes were also probed with anti-Actin antibodies or stained with Ponceau S for Rubisco subunit to indicate comparable loading between extracts. Migration position of protein molecular weight standards (in kDa) are indicated.

### Enhanced defenses in *sum1-1* is SA-dependent

To genetically determine whether constitutive SA-signaling routes impart increased defenses to *sum1-1*, we generated *sum1-1sid2-1* and *sum1-1eds1-2* double mutants. A *sid2-1* plant harbors a null mutation in *SID2/ICS1* whereas an *eds1-2* expresses a non-functional EDS1 (ENHANCED DISEASE SUSCEPTIBILITY1), a central player orchestrating SA-mediated defenses [43,49]. In segregating F2 populations, we noted that phenotypic defects always associated with homozygous *sum1-1* allele provided at least one functional copy of *EDS1* or *SID2* were present (data not shown). Remarkably, the reduced leaf mass apparent in *sum1-1* was improved in *sum1-1eds1-2* and *sum1-1sid2-1* plants (Fig 5A; S1 Fig). While *sum1-1eds1-2* achieved wild-type mass, abolishing *SID2/ICS1* in *sum1-1* although ameliorated growth defects but these plants still retained *sid2-1*-characterstics including smaller and paler leaves than Col-0 (Fig 5A; S1 Fig). Interestingly, both *sum1-1sid2-1* or *sum1-1eds1-2* had wild-type trichome densities unlike the *sum1-1* parent thus associating these defects to perturbations in SA-regulated networks (S6A Fig).

**Fig 5:**
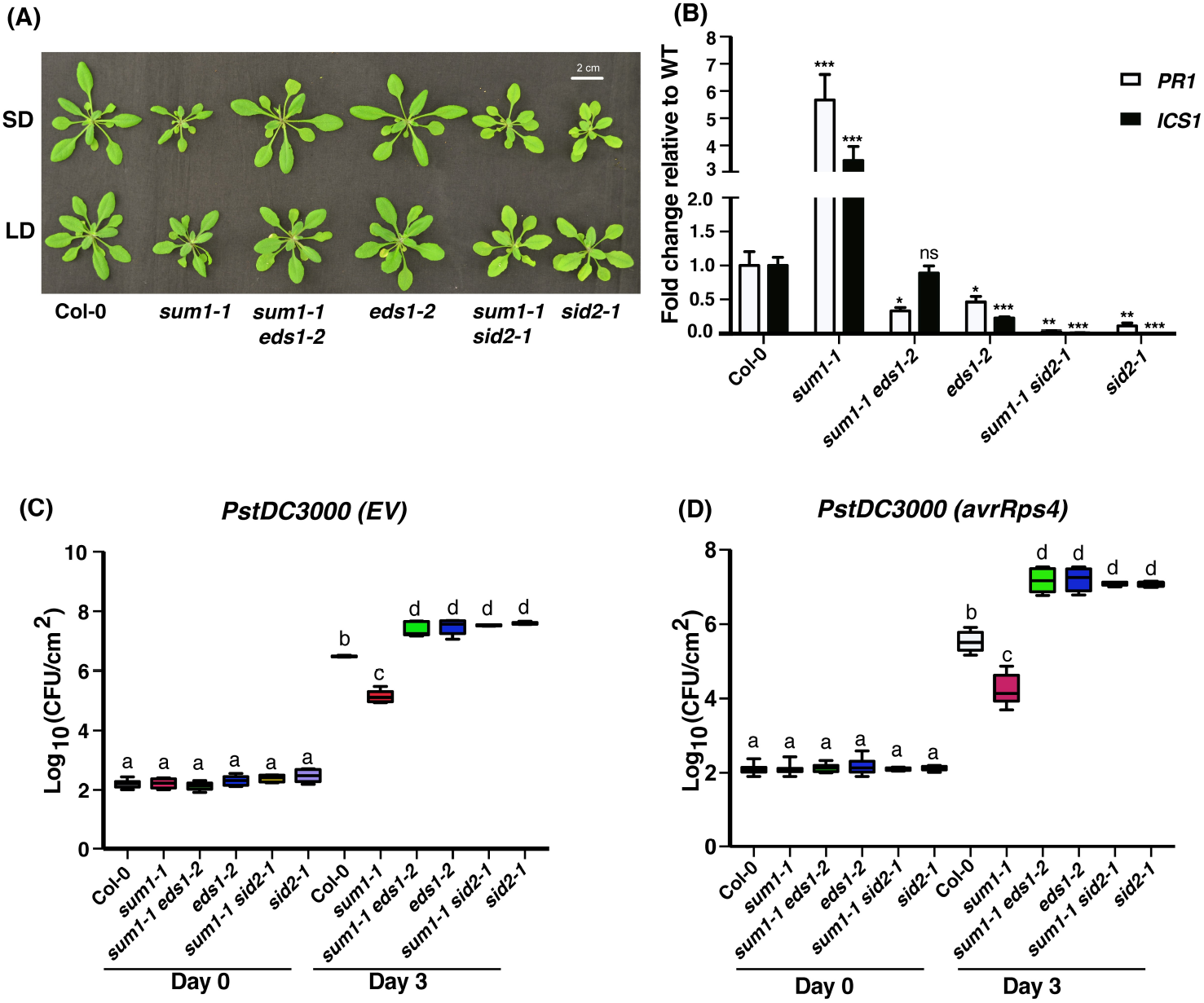
Growth retardation and enhanced defenses in *sum1-1* are SA-dependent. (A) Developmental phenotypes of 4-week-old Col-0, *sum1-1*, *sum1-1 eds1-2*, *eds1-2*, *sum1-1 sid2-1*, and *sid2-1* plants grown under SD and LD growth conditions. (B) Relative transcript abundance of *PR1* and *ICS1* in 4-week-old SD grown indicated plants was determined by qRT-PCR and normalized to *SAND* expression. The values are represented as fold change relative to Col-0 (WT). Data is representative of independent experiments (n=3). Error bars indicate SD. Student’s t-test significances are mentioned (*=*p*<0.05; **=*p*<0.01; ***=*p*<0.001). Growth of (C) virulent *PstDC3000(EV)* and (D) avirulent *PstDC3000* (*avrRps4*) strains, respectively in indicated plants are shown. Three to four expanded leaves from 4-week-old SD grown plants of each genotype were infiltrated with the indicated bacterial suspension at a density of 5 × 10^4^ cfu ml^−1^. Leaf discs were punched from the infiltrated area, macerated in 10mM MgCl_2_, serially diluted and plated on appropriate antibiotic plates. Bacterial titer was calculated at 0- and 3-dpi for the indicated infiltration. Whisker Box plot with Tukey test; n= 12-18; ANOVA was performed to measure statistical significant differences (at *p-value* <0.001) in growth of bacteria in Log_10_ scale.

Loss of either *EDS1* or *SID2*/*ICS1* remarkably abolished accumulated PR2 proteins and upregulated *PR1*, *PR2*, *FRK1* and *WRKY29* transcripts in *sum1-1* (Figs 4A, 5B; S6B Fig). Further on, pathogen growth assays using virulent and avirulent *PstDC3000* strains demonstrated that *sid2-1* or *eds1-2* mutations were epistatic to *sum1-1* abolishing not only the increased defenses but also conferring hyper-susceptibility to the respective double mutants towards virulent *PstDC3000(EV)* or avirulent (*avrRps4*- or *hopA1*-expressing) strains (Figs 5C,D; S6C Fig). Neither *eds1-2* nor *sid2-1* mutation affected resistance towards avirulent *PstDC3000* (*avrRpm1*) which remained comparable to *sum1-1* or Col-0 (S6D Fig). As expected, loss of *EDS1* or *SID2/ICS1* prevented accumulation of PR1 or PR2 proteins during *PstDC3000* infections (Figs 4B,C). With these assays, we identify that enhanced resistance in *sum1-1* is undoubtedly due to constitutive SA-routed defense signaling. A *sum2-1eds1-2* double mutant, we also generated, mimicked *sum1-1eds1-2* hyper-susceptible outcomes in disease assays (S7 Fig). Overall, our results provide several parallel lines of evidences suggesting *SUM1/2* intersections on SA-mediated immune signaling.

### *sum3-1* alleviates enhanced defenses in *sum1/2* mutants

Since our data suggested antagonistic involvement of *SUM1/2* and *SUM3* as negative or positive defense regulators, respectively we surmise a crosstalk between these isoforms. To investigate this, we generated *sum1-1sum3-1* double mutant by genetic crossing. Unlike *sum1-1eds1-2* or *sum1-1sid2-1*, only partial restoration of *sum1-1* growth defects was observed in *sum1-1sum3-1* plants (Fig 6A; S8A,B Figs). These observations hinted that *SUM3* is mildly responsible for *sum1-1* growth anomalies. Increased trichome densities noticed in *sum1-1* however remained unaffected by the loss of *SUM3* (S6A Fig). Because *SUM3* affects basal SA accumulation (Figs 2A,B), we tested defense outputs are altered in *sum1-1 sum3-1*. Elevated SA levels in *sum1-1* were marginally reduced in *sum1-1 sum3-1* plants. Indeed, the expression of *SID2/ICS1* remained comparable between *sum1-1* and *sum1-1 sum3-1* plants (Fig 2C). Additionally, *PR1* or *PR2* transcripts in *sum1-1 sum3-1*, were intermediate between *sum1-1* and Col-0. These data imply *SUM3* does not affect SA-biosynthesis but modulates positive feedback loop of SA-signaling wherein it intersects with *SUM1* role as a transcriptional repressor of *SID2/ICS1*, *PR1* or *PR2*. As is therefore expected, loss of *SUM3* reduced PR1 or PR2 levels in *sum1-1* (Fig 4A). Not the least, we also demonstrate that increased SNC1 accumulation in *sum1-1* plants although reduced in *sum1-1 sum3-1*, are still maintained higher than Col-0 levels (Fig 6B).

**Fig 6:**
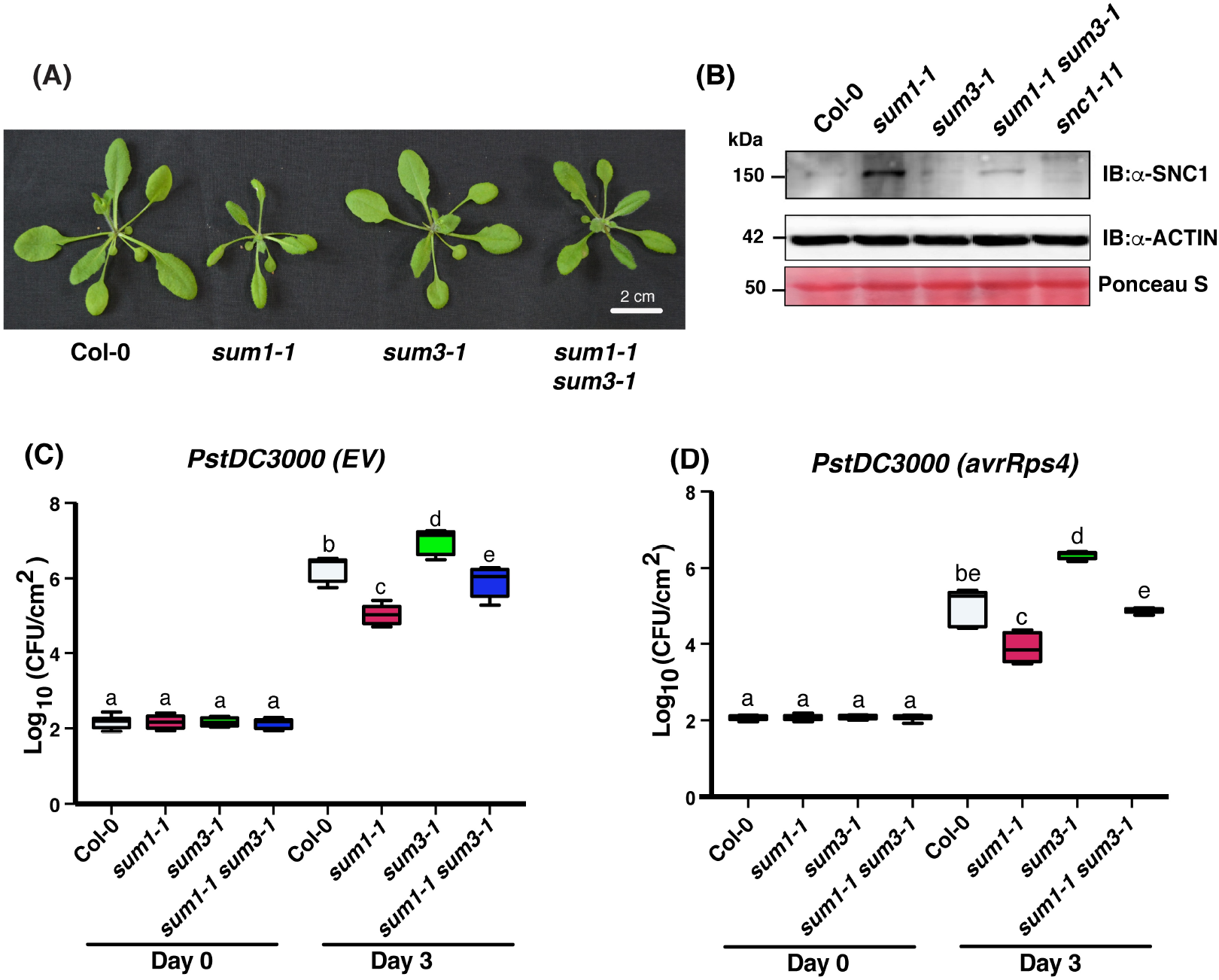
*sum3-1* alleviates developmental defects and enhanced defense responses in *sum1-1*. (A) Developmental phenotypes of 4-week-old SD grown Col-0, *sum1-1, sum3-1* and *sum1-1 sum3-1* plants. (B) Total protein extracts from indicated plants were immunoblotted with anti-SNC1 antibodies. Ponceau S stain of the Rubisco subunit and anti-Actin immunoblot indicative of equal protein loading between samples are shown. Migration positions of molecular weight standards (in kDa) are indicated. Growth of (C) virulent *PstDC3000(EV)* or (D) avirulent *PstDC3000(avrRps4)* strains, respectively in indicated plants. Three expanded leaves from 4-week-old SD grown plants of each genotype were infiltrated with the respective bacterial suspension at a density of 5 × 10^4^ cfu ml^−1^. Leaf discs of predefined diameter were punched from the infiltrated area, macerated in 10mM MgCl_2_, serially diluted and plated on appropriate antibiotic plates. Bacterial titer was calculated at 0- and 3-dpi with for indicated infiltration. Whisker Box plot with Tukey test; n= 12; ANOVA was performed to measure statistical significance (at *p-value* <0.001) in growth of bacteria in Log_10_ scale.

In pathogen growth assays, either virulent *PstDC3000(EV)* or avirulent *PstDC3000* (*hopA1*) strains accumulated to intermediate levels in *sum1-1 sum3-1* leaves, particularly higher than *sum1-1*, but significantly lower than in Col-0 or *sum3-1* (Fig 6C; S8C Fig). Interestingly, wild-type level of resistance was observed in *sum1-1 sum3-1* to the avirulent effector *avrRps4* (Fig 6D). Growth of avirulent *PstDC3000* (*avrRpm1*) was not affected in *sum1-1sum3-1* and remained comparable in all plants (S8D Fig). As investigated earlier, in response to virulent and avirulent *PstDC3000(avrRps4)* infections and unlike *sum1-1*, *sum1-1 sum3-1* remained deficient in induction of PR1 or PR2 at 6-hpi (Figs 4B,C; S9A,B Figs). However, we observed that at later time points (12- and 24-hpi) loss of *SUM3* did not affect the rapid induction of PR1/PR2 in *sum1-1*. This is a sharp contrast to *sum3-1* plants which even at 24-hpi accumulated very less PR1/PR2 proteins. Thus, although *SUM3* is deemed essential for upregulation of PR1/PR2 upon a pathogen attack, *sum1-1* plants eventually overcome this requirement. Taken together, we principally support that *SUM3* promotes *PR1*/*PR2* transcription downstream of SA as suggested earlier [21]. We provide further molecular evidence that *SUM1* suppression of defenses likely is upstream of SA and may involve transcriptional repression of *SID2/ICS1* and its subsequent consequences on SA-responsive markers such as *PR1*, *PR2*, *FRK1*, *WRKY29* and *SNC1* among others.

Adjacent chromosomal arrangement of *SUM2* and *SUM3* in Col-0 genome suggest that they likely arose by a tandem duplication event [18]. Hence, unlike a *sum1-1 sum3-1* double mutant, a *sum2-1 sum3-1* is difficult to obtain by genetic crossing. To test whether *SUM3* also intersects on *SUM2* function as a negative immune regulator, we generated *amiR-SUM2 sum3-1* plants by crossing the two parental lines. The *amiR-SUM2* plants reported earlier contains specific knockdown of *SUM2* and mimics the null mutant *sum2-1* [21]. *SUM1* or *SUM3* transcripts remain unaffected in these lines. Plants homozygous for *sum3-1* and containing *amiR-SUM2* transgene have significantly downregulated *SID2/ICS1, PR1* and *PR2* levels in comparison to a *sum2* mutant (Fig 2C). These transcripts however are still elevated than Col-0 plants implicating that *SUM3* functions impinge partially on *SUM2* role as a transcriptional repressor. This was also supported in pathogen challenges wherein growth of both virulent and avirulent *PstDC3000* strains were intermediate between resistant Col-0 and hyper-resistant *sum2* mutant (S7B,C Figs). Taken together, whether the functional antagonism in immune regulatory roles of *SUM1/2* and *SUM3* affect immune outcomes in plants.

### SUMO3, but not SUMO1, can form non-covalent homo-dimers

Conjugation proficiencies *in planta* of SUMO isoforms are clearly different. Upon SA application, drastic increase in SUMO1/2 conjugates are observed whereas target modifications by SUMO3 is barely detectable even though *SUM3* expression, unlike *SUM1/2* is SA-inducible [6,21]. A distinct difference in the protein sequence of these isoforms is the presence of a predicted SIM motif in SUMO3 but not SUMO1/2 (S10 Fig). We utilized Bi-molecular Fluorescence Complementation (BiFC) assays to test homo- and hetero- non-covalent associations between SUMO1 and SUMO3. The BiFC vectors introduced via *Agrobacterium*-mediated transient transformation of *N. benthamiana* leaves expressed split YFP fusions of either SUMOylation-proficient (GG) or -deficient (AA) SUMOs wherein the C-terminal di-glycine (GG) residues were kept intact or mutated to di-alanine (AA), respectively (Fig 7A). Co-expression of only SUMO1GG/SUMO1GG, but not SUMO1GG/SUMO1AA or SUMO1AA/SUMO1AA showed reconstitution of the split YFP protein. Although positive BiFC indicate non-covalent interactions among protein partners, the lack of fluorescence in SUMO1AA/SUMO1AA suggests that SUMO1GG/SUMO1GG combinations and not homo-oligomers and likely reflect covalent polySUMO1-conjugates wherein the split fluorescent fusion proteins are in allowed proximity for reconstitution. Remarkably, all BiFC combinations of SUMO3 with itself, regardless of GG- or AA-forms, showed YFP fluorescence. Since SUMO3 lacks poly-SUMOylation properties [22], we reason that the observed associations are non-covalent SUMO3 oligomers. To test whether SUMO3 binds SUMO1 non-covalently, combinations of SUMO1GG or AA were co-expressed with SUMO3GG or AA. While SUMO1GG showed clear fluorescence when expressed with either SUMO3GG or AA, similar combinations of SUMO1AA did not. These observations are suggestive that only a conjugable-SUMO1 likely present on a target may achieve permissible molecular proximity *in vivo* to a SUMO3 that may be either bound to a SIM (as in SUMO1GG/SUMO3AA pair) or attached covalently to the same target at a separate SUMOylation motif (as in SUMO1GG/SUMO3GG pair). Evidences from human cell lines demonstrate that ortholog of *Arabidopsis* SUMO3, HsSUMO1 ‘cap’ poly-SUMO chains of the SUMO1 ortholog HsSUMO2 [50,51]. Whether *Arabidopsis* SUMO3 regulates poly-SUMO1/2 chain lengths remain a promising possibility to explore further. Since no interaction was detected with SUMO1AA/SUMO3AA, it is clear that SUMO3 and SUMO1 do not interact non-covalently. To validate further, we performed *in vitro* binding assays with tagged recombinant SUMO1AA/SUMO3AA proteins expressed in *E. coli.* Enrichment of His-SUMO1AA via Ni^2+^ beads failed to co-elute either Strep-SUMO1AA or Strep-SUMO3AA suggesting that SUMO1-SUMO1 homo- and SUMO1-SUMO3 hetero-dimers cannot form *in vitro* (Fig 7B). Homodimers of SUMO3AA however were detected by the presence of Strep-SUMO3AA in His-SUMO3AA-enriched eluates. These results support our hypothesis that BiFC interactions observed for SUMO1GG with SUMO3GG/AA may likely suggest close proximity, but not direct binding, of SUMO1 and SUMO3 *in planta*.

**Fig 7:**
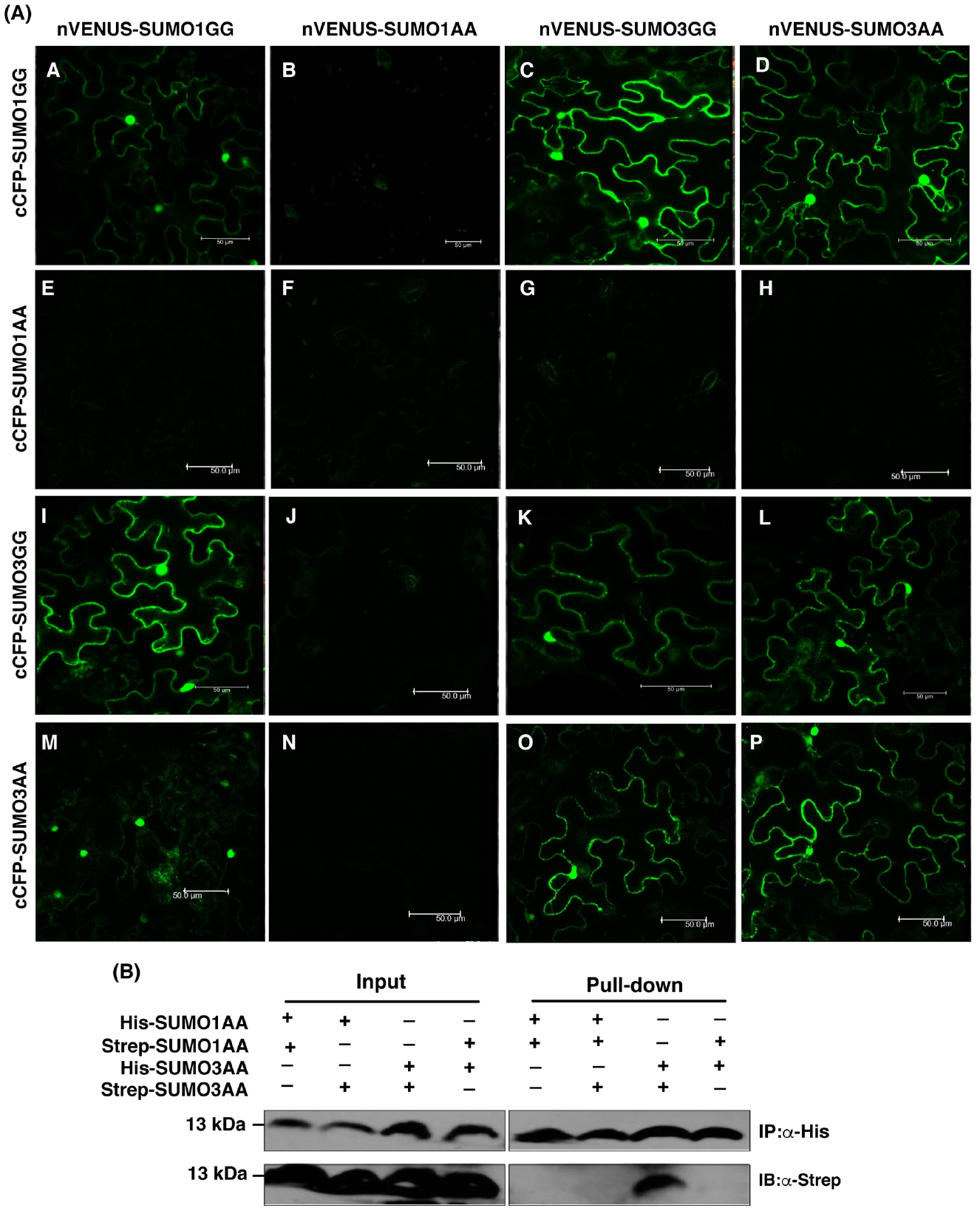
SUMO3, but not SUMO1, can form oligomers. (A) Green fluorescence indicative of YFP reconstitution between BiFC (Bi-molecular Fluorescence complementation) combinations of SUMOylation proficient (GG) or deficient (AA) forms of SUMO1 and SUMO3 isoforms are shown. *Agrobacterium* GV3101 strains expressing the indicated BiFC constructs were co-infiltrated in expanded *N. benthamiana* leaves. Infiltrated leaf sections were visualized under a confocal microscope at 2-days post-infiltration (dpi). (Scale bar = 50 μm). Images are representative of pattern observed in two independent experiments. (B) *In vitro* binding assays between SUMO1 and SUMO3 isoforms. His-or Strep-II-tagged SUMO1 or SUMO3 isoform as indicated were expressed in *E. coli.* Bacterial lysates from indicated combinations were mixed and enriched through the Ni^2+^-NTA matrix. Enrichment of the mentioned His-SUMO and the co-eluting isoform was investigated via immunoblotting with anti-His or anti-Strep antibodies, respectively (pull-down panel). The input panel shows the protein levels in the extracts used for the enrichments. Position of protein molecular weight standards (in kDa) are indicated.

### *SUMO3* regulates *SUMO1/2* conjugation efficiencies upon various stress exposures

With the above results, it is encouraging to speculate that SUMOylation efficiencies may be modulated by intersecting SUMO1-SUMO3 functions. Since *SUM3* partially modulates *sum1-1* defenses, we investigated dynamic changes in SUMO1/2-SUMOylome in Col-0, *sum1-1*, *sum3-1*, or *sum1-1 sum3-1* upon a pathogen challenge. Leaf tissues either mock-treated or infiltrated with *PstDC3000*(*EV*), were collected at 24-hpi and processed for selective qPCRs or immunoblots for SUMO1/2-conjugates. The polyclonal anti-SUMO1 antibody we used cannot distinguish SUMO1/2 isoforms and the lack of anti-SUMO3 antibodies prevented us for analyzing SUMO3-conjugates. We observed that while a pathogen exposure caused remarkable increase in SUMO1/2-conjugates in Col-0, for *sum1-1* these was considerably lower (Fig 8A).

**Fig 8:**
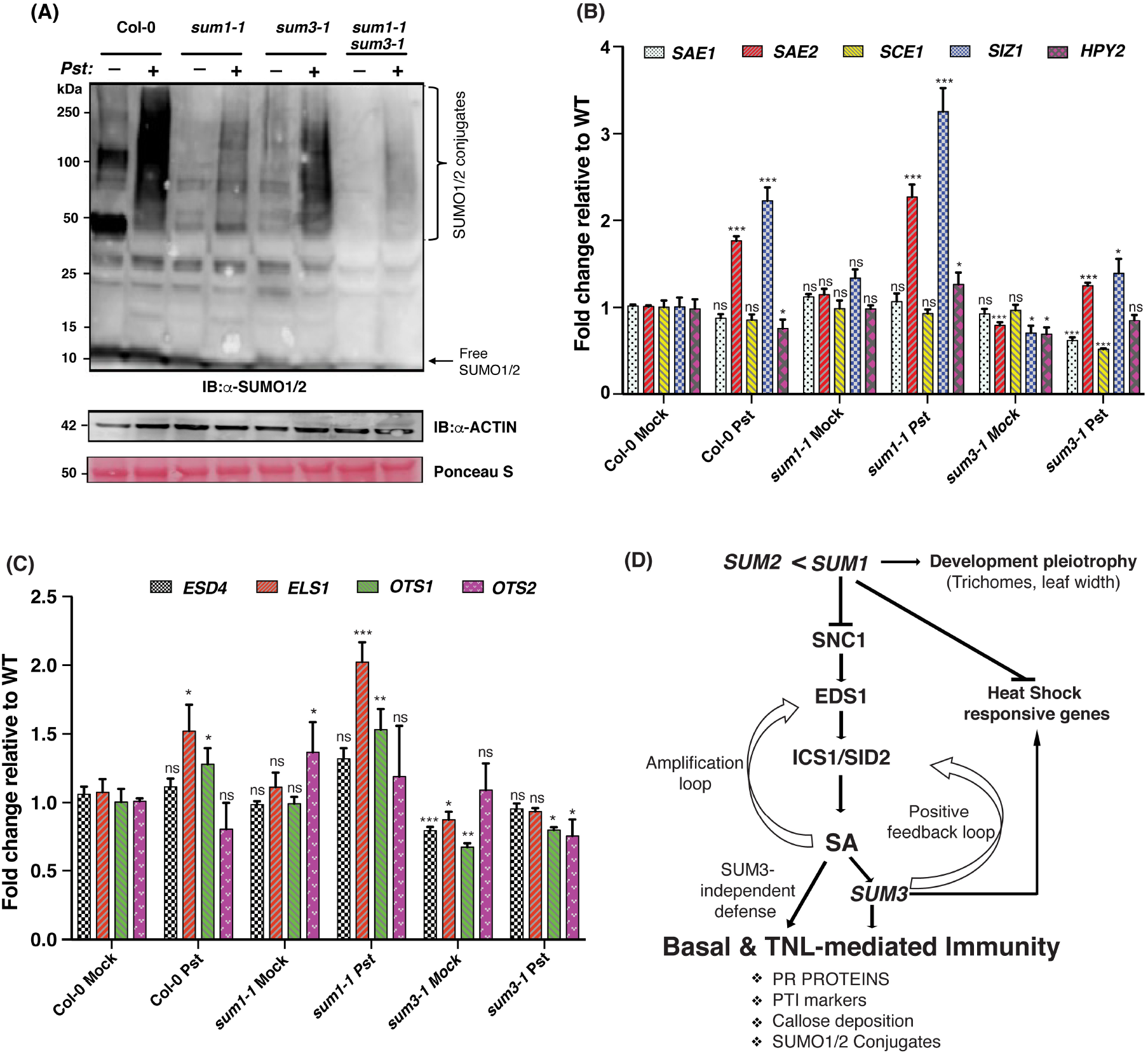
Loss of *SUM3* reduces SUMO1/2-conjugate enhancements in response to *PstDC3000(EV)* infection. (A) Fully expanded 3-4 leaves from 4-week-old SD grown Col-0, *sum1-1, sum3-1* and *sum1-1 sum3-1* plants were infiltrated either with buffer alone (-lanes) or with the virulent *PstDC3000(EV)* strain (+ lanes) at a density of 5 × 10^6^ cfu ml^−1^. Total protein isolated from the infected leaves at 24 hrs post-infiltration (hpi) was used for immunoblotting with anti-SUMO1/2 antibodies. Approximate positions of SUMO1/2-conjugates are shown. Anti-Actin immunoblot or Ponceau S straining demonstrate comparable protein loadings from the extracts. Relative positions of protein molecular weight standards (in kDa) are indicated. Relative transcript abundance at 24-hpi of (B) *SAE1, SAE2, SCE1, SIZ1, HPY2* and (C) *ESD4, ELS1, OTS1, OTS2* in mock versus *PstDC3000(EV)* infiltrated samples of Col-0, *sum1-1* and *sum3-1* was determined by qRT-PCR and normalized to *SAND* expression. The values are represented as fold change relative to Col-0 (WT). Data is representative of mean of three biological replicates (n=3). Error bars indicate SD. Student’s t-test was performed to calculate statistical significance; *=*p*<0.05; **=*p*<0.01; ***=*p*<0.001, ns= not significant. (D) Genetic model for *SUM1/2* crosstalks with *SUM3* in regulation of SA-dependent defences and heat-shock responses in *Arabidopsis*. *SUM1/2* function additively but not equally as negative regulators of defences via modulation of SA-signaling routes and expression of defense-associated markers. Pathogen exposure induce SA-dependent *SUM3* role as a positive regulator leading to increase in SUMO1/2-conjugation proficiencies, and expression of response-appropriate markers such as PR proteins, PTI markers, and callose deposits. Induced SUMO1/2-conjugates modulate SUMO3 functions and prevents overshooting of responses. *SUM1* also regulates via *SUM3*-independent but SA-dependent developmental aspects such as trichome production and leaf textures.

These results suggested that SUMO1-rather than SUMO2-conjugates are not only more prevalent but also undergo massive increment upon pathogen treatment. Surprisingly, the conjugates are also distinctly lower in uninfected *sum3-1* and are considerably less upregulated than Col-0 in response to the pathogen infection. As is expected, in *sum1-1 sum3-1* extracts barely any SUMO-conjugates are detectable pre- or post-infection. With these results, we identify clear intersection of SUMO3 functions on SUMO1/2-conjugation efficiencies and its perturbations during a pathogen attack. We also observed that both *sum1-1:His-H89R-SUM1* and *sum3-1:HS-SUM3* plants achieved Col-0 levels of SUMO1/2-cojugates upon *PstDC3000*(*EV*) infection thus further validating the complete functional complementation of *sum1-1* or *sum3-1*, respectively (S11 Fig).

To determine whether changes in conjugation efficiencies are due to differential expression of SUMO isoforms and/or SUMOylation-associated genes, we investigated their relative expressions in Col-0 and the *sum* mutants (Figs 8B,C; S11B Fig). We observed that *SUM3*, but not *SUM1/2*, was upregulated upon a *PstDC3000*(*EV*) challenge. This is in accordance with *SUM3* being SA-inducible [21]. Additionally, we noticed that the enhanced *SUM3* expression in *sum1-1* was further boosted in pathogen-exposed samples. Curiously, for the investigated SUMOylation-associated genes although *sum1-1* mutation did not significantly alter their basal expression levels, several of these (*SAE2*, *SIZ1*, *HYP2*, *ESD4*, *ELS1*, and *OTS1*) were down-regulated in *sum3-1* plants. Thus, the modest reduction in global SUMOylome in *sum3-1* we observe may be attributed to *SUM3*-dependent modulation of expression of these genes. Interestingly, *sum1-1 sum3-1* plants showed slight upregulation of *SAE2*, *SCE1*, *SIZ1*, *HPY2*, *ELS1* and *OTS2* in mock-treatment supporting our earlier claim that *sum1-1* plants with constitutively active defenses and elevated SA levels, overcome the requirement of *SUM3* for the expression of these genes. We also noted that upregulation of *SAE2*, *SIZ1*, *ELS1* or *OTS1* that occurred only modestly in Col-0 upon *PstDC3000*(*EV*) infection were further aggravated in *sum1-1*, but intermediate in *sum3-1* (Figs 8B,C). These enhancements are maintained in *sum1-1 sum3-1* plants and are likely *SUM3*-independent. Remarkably, *sum3-1* plants down-regulated the expression of *SUM1* and *SUM2* only upon pathogen-treatment highlighting that the positive immune function of *SUM3* may involve transcriptional repression of negative defense regulators such as *SUM1/2.* Overall, our data lead us to speculate that immune responses recruit *SUM1*-*SUM3* crosstalks to modulate the expression of these genes at a transcriptional level thereby influencing host SUMOylome changes.

SAE2, SCE1, SIZ1 or EDS4 are direct SUMO1 targets and both SUMO1/3 interact *in planta* with SCE1 and SIZ1 [5,52]. These led us to test whether SUMO1-SUMO3 intersections directly influence the efficiency of SUMO1/2-conjugate formation through modulation of SUMOylation-associated protein functions. We utilized the *E. coli* SIZ1-independent *Arabidopsis* SUMOylation reconstitution system [53]. Co-expression of His-SUMO1, and Strep-SUMO3 (GG or AA) along with SCE1, and SAE1/2 showed modest but clear increase in SUMO1-conjugates in comparison to extracts that lacked Strep-SUMO3 (S12 Fig). Remarkably, a more dramatic increase in SUMO3-conjugates when co-expressed with either Strep-SUMO1 GG or AA was also noted. From these results, it is undeniably clear that SUMO1 and SUMO3 have a mutually beneficial role in reciprocal SUMOylation process likely through modulation of SCE1 or SAE1/2 functions. The significant decrease in SUMO1/2 conjugates in absence of *SUM3* (Fig 8A) provides reasonable endorsement to our hypothesis.

SUMO1/2-conjugates are enhanced in response to heat shock [20,21]. To test whether *SUM3* role intersect on these responses, we subjected Col-0, *sum1-1*, *sum3-1*, or *sum1-1 sum3-1* plants to heat shock and analyzed SUMO1/2-conjugates as well as relative expression levels of *SUM1/2/3* and several previously characterized heat stress-responsive genes [54] (S13 Fig). In accordance with [20], we observe strong induction of SUMO1/2-conjugates in heat-treated Col-0 extracts (S13A Fig). These conjugates were low in heat-stressed *sum1-1* plants reinstating that SUMO1-mediated SUMOylation is most predominant during heat stress. Surprisingly, in *sum3-1* considerably reduced conjugates than Col-0 accumulated upon heat shock. We especially noted that unlike *PstDC3000* challenges, *SUM1/2* transcripts were significantly upregulated by heat-exposure, while *SUM3* induction was almost negligible (S13B Fig). Further on, transcriptional upregulation of *SUM1/2* was *SUM3*-independent. Therefore, reduced accumulation of SUMO1/2-conjugates in heat-treated *sum3-1* suggests that *SUM3* affects SUMO1-SUMOylation efficiencies at a post-transcriptional level. With these results, we infer that although different physiological stresses may cause apparently similar effects on global SUMO1/2-conjugates, the responsive routes to condition these are not only distinct but also stress-specific. Interestingly, increased expression of several heat stress-responsive markers such as *HsfA2*, *HSP22.0-ER* that are upregulated in Col-0 upon heat-treatment, were hyper-elevated in *sum1-1* suggesting *SUM1* suppresses their expression (S13C Fig). In *sum3-1* plants, their fold-induction upon heat stress was relatively lower than Col-0 implying that *SUM3* is partially responsible for their upregulation. Induction of other tested markers such as *HSP18.2* or *HSP23.5-P* were unaffected in *sum1-1* or *sum3-1* plants (S13D Fig). Taken together, our results clearly identify *SUM1-SUM3* crosstalks in responses to multiple stresses.

## Discussion

Functional overlaps between *Arabidopsis* SUMO1 and SUMO2 isoforms, likely prevent noticeable phenotypic defects in the individual mutants under non-stressed conditions [17,20,21,23]. Interestingly, reduction in SUMO-conjugates both basally as well as upon heat-stress is more prominent in *sum1-1* than *sum2-1* suggesting a *SUM1* predominance in these responses [20]. Our observations therefore of phenotypic defects in *sum1-1* but not *sum2-1* plants may reflect a similar developmental process preferentially regulated by *SUM1*. We convincingly establish that constitutive upregulation of SA-regulated networks is the primary cause of these defects in *sum1-1* (S6A Fig). This is unlike *siz1-2* where the associated growth abnormalities are EDS1- or SID2/ICS1-independent [55]. We deduce that reduction in global SUMO1/2-conjugates apparently similar between *siz1-2* and *sum1-1* affect downstream responses differently. At one instance, loss of *SIZ1* may affect SUMO-conjugation for all isoforms, whereas the *sum1-1* clearly restricts only SUMO1 functions. Previous studies propose SA antagonism or jasmonic acid (JA) promotion in increased trichome formation [56]. We reason that while direct application of SA or JA may affect trichome production as reported, the loss of *SUM1* whose substrates include both JA and SA signaling regulators such as SIZ1, TPL, or JAZs among others may impact trichome density through a more complex SA-JA crosstalk [5,57]. Further investigations into relative SA-JA signaling routes perturbed in *sum1-1* may unravel this mystery.

In context of defense responses, our data provide substantial molecular support not only to individual *SUM1/2* or *SUM3* but also to their intersecting contributions as negative or positive regulators, respectively of anti-bacterial basal and TNL-specific immunity. Firstly, bacterial accumulations are strongly reduced in both *sum1-1* and *sum2-1*, with *sum1-1* displaying stronger immunity than *sum2-1*, for both virulent as well as avirulent *PstDC3000*(*hopA1*) infections. Although comparable immunity between *sum1-1* and *sum2-1* is observed for avirulent *PstDC3000*(*avrRps4*) challenges, this likely is due to relatively weaker ETI responses elicited by *AvrRps4* in comparison to *HopA1*-expresssing *PstDC3000* (compare Fig 1D and S2A Fig). In contrast to *sum1/2* mutants, immunity in *sum3-1* is compromised implicating *SUM3* is essential for these defenses. Secondly, we demonstrate that impairment of SA-signaling routes due to loss of *EDS1* (*eds1-2*) or *SID2/ICS1* (*sid2-1*) abolish enhanced resistance in *sum1-1* and *sum2-1* plants providing support to genetic placement of *SUM1/2* as suppressors of these defenses [21] (Fig 8D). In this pathway, *SUM3* partially delegates SA-defenses likely through positive feedback mechanisms. Indeed, both *sum1-1* and *sum2-1* plants accumulate increased SA than Col-0, while in *sum3-1*, these are relatively lower. Correspondingly, transcripts of SA-biosynthesis *SID2/ICS1* and responsive markers such as *FRK1*, *PR1*, *WRKY29*, and *SNC1* are upregulated in *sum1-1* or *sum2-1*, and lower in *sum3-1*. Accumulation kinetics of PR1/PR2 proteins or transcripts (Fig 4; S5 Fig) further validate primed or deficient SA-mediated defenses in *sum1-1* or *sum2-1* and *sum3-1*, respectively. The modest difference we note between *sum1-1* and *sum2-1*, in the upregulation of some of these SA-inducible markers, is suggestive of additive but unequal roles of *SUM1* and *SUM2* as negative immune regulators. Only ~17% common targets were identified between human SUMO1 and SUMO2 isoforms suggesting that they are partially redundant at best [58]. Whether SUMOylation-targets distinct between *Arabidopsis* SUMO1 and SUMO2 isoforms affect immune amplitudes differentially awaits further studies.

We provide the first *in planta* evidence of partial SUMO3 moderation on SUMO1/2 functions in biotic, abiotic and developmental responses. Introducing *sum3-1* mutation not only attenuates reduced tissue mass but also subdues enhanced resistance of *sum1-1* supporting *SUM3* role in promoting defenses downstream of SA. Thus, elevated SA and *PR1*, *PR2*, or *SNC1* expressions are partially dampened in either *sum1-1* or in *amiR-SUM2* plants when *SUM3* is mutated (Fig 2). Intersections of *SUM3* functions on SUMOylation-proficiencies, are also best noted on SUMO1/2-conjugate intensities in response to different stresses. Although, enhancement upon heat shock or SA-treatments have been previously reported [5,6], we first report their upregulation in *PstDC3000* exposures. Induction of several SUMOylation-associated genes such as *SAE2*, *SIZ1*, *ELS1*, or *OTS1* during *PstDC3000* challenges are influenced by *SUM1*-*SUM3* intersections and contribute to the corresponding defense outcomes (Fig 8). Recently, the activity of TPR1, a transcriptional co-repressor involved in suppression of negative defense regulators *DND1/2* was reported to be regulated in a partial SA-dependent manner via SUMOylation by SIZ1 [59]. SUMOylation-deficient TPR1 not only is more enhanced in repressing *DND1/2* expressions, but also its over-expression cause enhanced resistance than the SUMOylation-proficient TPR1. Overall, our results raised the possibility that increased SUMO1/2 conjugates may reflect negative feedback mechanisms to maintain immune responses as transitory thus regulating *SUM3* functions in promoting defenses.

Functional intersections among SUMO isoforms are anticipated at multiple post-transcriptional events. Amino acid distinctions at key conserved positions among the SUMO isoforms influence their *in vivo* conjugation proficiencies [60] (Schematically summarized in S10 Fig). Aspartate63 (D^63^) in *Arabidopsis* SUMO1 (D^75^ in SUMO2) is replaced by Asparagine63 (N^63^) in SUMO3, reducing its relative interaction strength than SUMO1 with SCE1, thus lowering its poly-SUMO formation efficiencies. Other residues also diverged in SUMO3 weaken thioester-bond formation and interaction with SAE1. Whether *SUM3* upregulation during defences improve these propensities and hence alters host SUMOylome outcomes although remains unknown, evidential support is obtained from several studies. Mutants with decreased SUMOylation including *siz1-2* have increased SCE1 protein abundance [15,20,61]. And *siz1-2* impairment in accumulating SUMO-conjugates during heat stress, is completely recovered in *pial1 pial2 siz1-2* plants [8]. Considering PIAL1/2 improves SCE1 ability to form otherwise less efficient polySUMO3 chains, defense responses with upregulated *SUM3* likely introduces competition between SUMO1 and SUMO3 for substrate polySUMOylation thereby altering their fates via respective isoform-specific SUMO-targeted E3 Ubiquitin ligases (STUbLs) [8].

Indeed, *pial1/2* double mutants although less stress-tolerant, have upregulated levels of proteins related to biotic stress [8,15]. In *sum1-1*, the reduced global SUMO1/2-conjugates upon pathogen or heat-shock challenges taken together with elevated *SUM3* transcripts is suggestive of this occurrence and need to be explored further.

As substrates of Arabidopsis SCE1 and ESD4, candidates predominantly involved in RNA-related processes such as nucleocytoplasmic transport, splicing, or turnover and chromatin-modification including transcriptional activation/repression-associated proteins were identified [62]. A majority of these candidates possessed predicted SUMOylation motifs and were covalently modified by both SUMO1 or SUMO3. In *Arabidopsis*, a SUMOylation-proficient SUMO1 is essential to covalently charge SCE1 at the catalytic site, promote its association with SIZ1, and form the ternary complex (SUMO-SCE1-SIZ1) [52]. Interestingly, the subcellular localization of this complex is determined by the specific SUMO isoform bound non-covalently at the SIM site distinct from the catalytic pocket of SCE1. Since SCE1 possess ability to distinguish substrates for SUMOylation [9], selection of substrates and formation of polySUMO chains therefore may be affected by fluctuations in relative SUMO1/3 levels, as has been previously suggested in animal studies [63,64]. Our data showing reciprocal improvements in SUMOylation efficiencies *in vitro* regardless of conjugation-proficiencies of the influencing SUMO isoform may suggest towards one such consequence of SUMO1-SUMO3 crosstalks (S12 Fig). Likewise, constitutive activation of defenses due to over-expression of either conjugation-proficient (GG) or -deficient (ΔGG) SUMO isoforms may at least be partially attributed to this phenomenon [21].

SUMOylation machineries such as SAE2, SIZ1 and ESD4 also bind SUMOs non-covalently owing to their intrinsic SIMs [12,65]. It is postulated that spatio-temporal regulations of SUMOylation/de-SUMOylation coordination primarily by SIZ1 and ESD4 regulate steady-state levels of host proteins conjugated to SUMOs which surprisingly are fewer in numbers [2]. Dramatic changes in this minimal SUMOylome across various stress conditions reveal that instead of newer targets undergoing covalent modifications by SUMOs, SUMOylation levels on prior-SUMOylated proteins pools are more altered [66]. Likely, functional inactivation of SUMO proteases especially noted post-stress in mammalian systems [67–69] and enhanced pool of SUMOylated pool SIZ1 [66] coordinate to achieve this. Taking this into account, a host SUMOylome output is likely to be influenced by relative changes in SUMO isoforms. Indeed, we demonstrate that transcripts of *SIZ1*, *EDS4* or *SAE2* are up-regulated basally in *sum1-1*, and in both Col-0 or *sum1-1* upon pathogen challenge in a *SUM3*-dependent manner (Fig 8). Hence the requirement of *SUM3* to maintain and adjust optimal level of SUMO1/2 SUMOylome is clearly evident. Non-covalent associations with SUMOs also affect chromatin architecture via their interaction with DNA methyltransferases and demethylases [11]. Expression of FLOWERING LOCUS C (FLC), a central transcription factor of flowering is dynamically regulated by DNA methylation status while the protein function is modulated by HPY2, SIZ1 and SUMO proteases [70,71]. FLC regulation presents a classical example of this versatility of SUMO-influences on a developmental process.

Stimulus-driven SUMO isoform switches and subsequent fate of substrates have been widely documented in animal systems [37–39]. The mammalian GTPase activating protein RanGAP1 although is equally modified by SUMO1/2/3 *in vitro*, conjugation *in vivo* to SUMO1, but not SUMO2, imparts more stability from isopeptidases thereby facilitating its association with Nup358 [39]. Similarly, HDAC1 is targeted for degradation upon SUMOylation by SUMO1, but not SUMO2, in cancerous cell lines thus potentiating invasive properties of the tumour [72]. Specific SUMO-proteases also regulate stimulus-dependent SUMO isoform switching [73].

Upon arsenic exposure, SUMO2 at Lys^65^ of PML (Promyelocytic leukemia protein) is replaced with SUMO1 to direct its ubiquitinylation. Clearly with the isoform selectivity regulated at multiple levels, a host SUMOylome undergoes dynamic changes in response to physiological perturbations. Evidences on SUMO isoform switches however are completely lacking from plant systems. Nonetheless several reports suggest the likelihood of this phenomenon. The effector XopD from *Xcv* is a SUMO3-specific isopeptidase [17]. The replication protein AL1 from Tomato Golden Mosaic Virus (TGMV) and the RNA-dependent RNA polymerase NIb from Turnip mosaic virus (TuMV) interact with and require SCE1 for replication [74,75].

While our results undoubtedly place SUMO3 as a strong contributor in regulating dynamics changes in a host SUMOylome, the loss of *SUM3* in many *Brassicaeae* and other higher eudicots still remain an enigma [18]. Whether functions of *SUM3* has been incorporated in *SUM1/2* roles in these plants remains to be explored further. We foresee the immediate need of a plant system that would facilitate enrichment via distinct affinity tags on individual SUMO isoforms in order to obtain evidences of SUMO preference, switches and intersections. To summarize, our data lay the foundation on functional impingements of *Arabidopsis* SUMO isoforms and their adjustments on global SUMOylome in response to physiological changes.

## Methods

### Plant materials and growth conditions

*Arabidopsis thaliana* mutant lines *sum1-1* (SAIL_296_C12), *sum2-1* (SALK_029775C) and *sum3-1* (SM_3_2707/SM_3_21645) were obtained from Arabidopsis stock centre (https://www.arabidopsis.org). Seeds of *amiR-SUM2 sum3-1* were generated by Dr. Harrold van den Burg. All plants were grown at 22°C with 70% humidity under Long Days (LD; 16 h: 8 h, light: dark) or Short Days (SD; 8 h: 16 h, light : dark) having light intensity 100 μmol μm^−2^s^−1^ light. Specific growth conditions are indicated in respective legends. To generate combinatorial mutants, *sum1-1* was genetically crossed with *eds1-2*, *sid2-1* or *sum3-1* mutant plants. The *sum2-1* mutant was similarly crossed with *eds1-2* to generate *sum2-1 eds1-2.* Double mutants (*sum1-1 eds1-2, sum1-1 sid2-1* and *sum1-1 sum3-1*) were identified in F2 population by PCR based genotyping. To generate *sum1-1:His-H89R-SUM1*, *His-H89R-SUM1* [5] was crossed with *sum1-1*. Segregating F2 populations were PCR-based genotyped for the presence of homozygous *sum1-1*, wild-type *SUM2* and homozygous copies of *His-H89R-SUM1* transgene. All primers used are listed in S1 Table.

### Trichome visualisation and quantification

For trichome visualization, leaves from 4-weeks old SD-grown plants were harvested and washed in Acetic acid: Ethanol (1:3) solution for overnight to bleach all pigments then visualised under bright field in a fluorescence microscope. The images were captured as 4848 × 3648 pixels from 4.1 mm^2^ leaf area. The trichome numbers were counted in 10 randomly selected images. Experiment was repeated twice with similar observations.

### Salicylic acid (SA) measurements

Free salicylic acid (SA) and glucose-conjugated SA (SAG) measurements were done using the *Acinetobacter* sp. ADPWH_*lux* biosensor system [76]. In brief, 100 mg leaf tissues were collected and frozen in liquid nitrogen. The tissue was homogenized in 250μl of acetate buffer (0.1 M, pH 5.6). Samples were centrifuged at 12000 rpm for 15 minutes. One aliquot (100μl) of the supernatant was used for free SA measurements, and another was incubated with 6U of β-glucosidase (Sigma-Aldrich) for 90 min at 37°C for total SA measurement. 20μl aliquot of each plant extract were added to 50μl of secondary culture of *Acinetobacter* sp. ADPWH_*lux* (OD600 of 0.4) with additional 60 μl of LB. Standard SA solutions (prepared in *sid2-1* extract) were also taken to generate standard curve to calculate the amount of SA present in samples. Plate was incubated at 37°C for 1 h, and luminescence detected using the POLARStar Omega Luminometer (BMG Labtech). Data shown is representative of three biological replicates with SD.

### *In planta* assays for bacterial growth and kinetics of defense-associated marker expressions

Bacterial growth assays were performed according to [46]. Briefly, *PstDC3000* strains were infiltrated with a needleless syringe at a density of 5 × 10^4^ cfu ml^−1^into fully expanded rosette leaves of 3-4-weeks-old SD grown plants. Leaf discs harvested from infiltrated area were macerated in 10mM MgCl_2_, serially diluted and plated onto selective medium plates. Bacterial growth was determined at 0 and 3-days post-infiltration. Bacterial infiltrations for kinetics of defensive markers were carried as above except a 5 × 10^6^ cfu ml^−1^ bacterial inoculum was used. Infiltrated samples harvested at 6, 12 and 24 hpi were processed separately for total RNA extraction and qPCR analysis or for immunoblots with indicated antibodies.

### Callose deposition assay and image analysis

The callose deposition assays were performed according to [77]. In brief, 4-week-old leaves of SD grown Col-0, *sum1-1, sum2-1* and *sum3-1* plants were infected with the indicated *PstDC3000* strains. At 24 hpi, 3-4 random infiltrated leaves were harvested and washed first in Acetic acid: Ethanol (1:3) solution for overnight to bleach all pigments and then washed for 30 min in 150 mM K_2_HPO_4_ solution. The leaves were then incubated in dark for 2 hours in 150 mM K_2_HPO_4_ containing 0.01% aniline blue in a 16-well tray. Samples were then embedded in 50% glycerol and observed under a Nikon fluorescence microscope with DAPI filter. Images were analysed using IMARIS 8.0 software for quantifying the number of callose deposits/ mm^2^ leaf area.

### RNA extraction and gene expression analysis by qRT-PCR

Total RNA extraction and cDNA synthesis was performed as described later. All qPCR primers used in this study are listed in S1 Table. qPCRs were performed in QuantStudio 6 Flex Real-Time PCR system (Applied Biosystems) with 5X HOT FIREPol^®^ EvaGreen^®^ qPCR Mix Plus (ROX) (Solis BioDyne) according to the manufacturer’s instructions. All qPCR experiments were replicated thrice with three biological and technical replicates (n=3). *SAND* (At2g28390) expression was used as internal control [46]. Relative expression was calculated according to the PCR efficiency^^−∆∆Ct^ formula. Expression differences were normalised to Col-0 and plotted as fold change.

### Protein extraction and western blotting

For immunoblotting, leaf tissues collected at indicated time points/treatments were homogenised in protein extraction buffer [50 mM Tris HCl (pH 8.0), 8M Urea, 50 mM NaCl, 1% v/v NP-40, 0.5% Sodium deoxycholate, 0.1% SDS and 1 mM EDTA] containing 20 mM *N*-ethylmaleimide (NEM), 1X plant protease inhibitors cocktail (Sigma Aldrich) and 2% w/v Polyvinylpyrrolidone (PVPP). The homogenates were clarified by centrifugation, mixed with 2X Laemmli buffer, proteins separated by SDS-PAGE and then transferred onto polyvinylidene fluoride (PVDF) membrane by wet transfer method. The membrane was blocked with 5% non-fat skim milk and western blots performed with indicated primary antibodies [anti-SNC1 (Abiocode), anti-PR1 or anti-PR2 (Agrisera), or anti-SUMO1 (Abcam), or anti-Actin C3 antibodies (Abiocode)] in 1X TBST at 4°C for overnight. Comparable protein loading was determined by Ponceau S staining of Rubisco subunit. Blots were washed thrice with 1X TBST and then incubated at RT for one hour with appropriate horse-radish peroxidase (HRP)-conjugated secondary antibodies. The blots developed using ECL™ Prime western blotting system (GE Healthcare) and visualised in ImageQuant™ LAS 4000 biomolecular imager (GE Healthcare).

### SUMOylome changes in response to SA or heat shock treatments

For salicylic acid (SA)-induced SUMOylation changes, extracts were obtained from 2-weeks old seedlings treated with 2 mM SA or buffer alone for 1 hour [6]. For heat shock treatments, 2-weeks old seedlings were incubated at 37°C or at RT for 30 minutes. Total protein extracts were immunoblotted as described earlier.

### Construction of Plasmid clones

To generate cDNA clones of Arabidopsis *SUM1* and *SUM3* genes, total RNA isolated from Col-0 plants (RNAiso Plus; Takara) was reverse transcribed (iScript™ cDNA Synthesis Kit; Bio-Rad) according to manufacturer’s instructions. Specific *SUM1* and *SUM3* (GG and AA forms) sequences were amplified (Phusion High-Fidelity DNA Polymerase; ThermoFisher) from the cDNA using the primers listed (S1 Table). PCR products were cloned into the Gateway entry vector *p*DONR201 and subsequently into *p*MDC-cCFP and *p*MDC-nVenus (BiFC destination vectors; [40] using the Clonase™ Recombination system (ThermoFisher). Confirmed BiFC clones were then electroporated into *A. tumefaciens* GV3101 strain.

For dimerization studies, cDNAs of *SUM1AA* or *SUM3AA* were cloned as EcoRI-SalI fragment into *p*ASK*-*IBA16 vector (N-terminal Strep-tag II affinity tag; Neuromics). For *in vitro* isoform influence on SUMOylation assays *SUM1GG* or *SUM3GG* were cloned as EcoRI-SalI fragment into *p*ASK*-*IBA16 vector. These clones were named accordingly as Strep-SUMO1GG, Strep-SUMO1AA, Strep-SUMO3GG, or Strep-SUMO3AA, respectively. For His-tagged versions, *p*CDFDuet vector harbouring *SUM1GG* (*p*KT-973), *SUM1AA* (*p*KT-1017), *SUM3GG* (*p*KT-975), or *SUM3AA* (*p*KT-1788) were used [53]. These plasmids were a kind gift from Prof. Katsunori Tanaka, Kwansei Gakuin University, Japan).

### Cloning and generation of *SUM3p-His-StrepII-SUM3g* transgenic plants

Genomic DNA extracted from Col-0 plants were used for PCR of a ~1.3kbp genomic fragment with KpnI-SUM3p For/XbaI-SUM3-UTR Rev primers. Using the amplicon as a template, two independent PCRs were performed by using KpnI-SUM3p For/His-StrepII Rev or His-StrepII For/ XbaI-SUM3-UTR Rev primers combinations respectively. Individual PCR amplifications were used for overlap PCR with KpnI-SUM3p For/XbaI-SUM3-UTR Rev primers. The product was cloned into XbaI-KpnI site of the binary vector *p*BIB-Hyg [78]. Generated clones were sequenced and confirmed. The binary vector was introduced in *Agrobacterium* strain GV3101 via electroporation. Pool of *sum3-1* plants were transformed via floral-dip method [79]. Transgenic *sum3-1:HS-SUM3* plants were selected on Hygromycin containing medium, propagated through T3 generations to identify lines containing single locus but homozygous *sum3-1:HS-SUM3* transgenes. Subsequently, the plants were used for assays as indicated.

### Bimolecular Fluorescence Complementation (BiFC) assays

All BiFC assays were performed according to [40]. Briefly, *Agrobacterium* cells harbouring the indicated BiFC vectors were induced with 150 μM Acetosyringone for 4 hours, equal bacterial density suspensions of desired cCFP and nVenus BiFC combinations made and then infiltrated in leaves of 4-weeks old *N. benthamiana* plants. At 48-hpi, a small section of infiltrated area was imaged under a SP8 Leica confocal microscopy system using FITC filter (488-nm Argon Laser).

### *In vitro* SUMO binding assays

SUMO-binding assays were performed with recombinant protein expressed in *E. coli.*

Expression was induced with 200 μg/L Anhydrotetracycline (for *p*ASK-IBA16 clones) or with 0.5 mM IPTG (for *p*CDFDuet clones) in BL21 (DE3) for overnight at 25°C. Cell pellet was resuspended in lysis buffer (50 mM NaH_2_PO4, 300 mM NaCl, 10 mM Imidazole; pH 8.0) and sonicated. Equal volumes of His- and Strep-II tagged-SUMO combination supernatants were mixed and incubated at 4°C with rotation for 2-3 hours to allow binding. Ni^2+^-NTA beads (Qiagen) were added to the mix and incubated further for 2 hrs. His-tagged protein pull down was performed under native conditions according to manufacturer’s instructions. Immunoblots were performed with anti-His-HRP (Santa Cruz Biotech) or anti-StrepII-HRP (IBA Lifesciences) antibodies.

### *In vitro* SUMOylation reconstitution assays

The His-tagged constructs used were according to [53]. Generation of Strep-tagged SUMO isoform clones have been described earlier. *E. coli* BL21 (DE3) cells containing both *p*KT-973 (SUMO1GG + SCE1) and *p*KT-978 (SAE2 + SAE1) was transformed with either empty vector (*p*ASK-IBA16), Strep-SUMO3GG, or Strep-SUMO3AA plasmids. Similarly, BL21 (DE3) cells containing *p*KT-975 (SUMO3GG + SCE1) and *p*KT-978 (SAE2 + SAE1) was either transformed with either empty vector (*p*ASK-IBA16), Strep-SUMO1GG, or Strep-SUMO1AA plasmids. As controls, conjugation-deficient SUMO1AA (*p*KT-1017) or SUMO3AA (*p*KT-1788) in combination with *p*KT-978 was used. Transformed cells were induced with 200 μg/L Anhydrotetracycline and 0.5 mM IPTG for overnight at 25°C. After harvesting, cell pellets were lysed and immunoblotted with anti-His or anti-Strep antibodies.

### Statistical Analysis

For all gene expression experiments, Student’s t-test was performed to check significance and denoted by one, two and three asterisks (*) indicating *p-value* <0.05, <0.01, and 0.001, respectively. For growth curve and other assays, ANOVA was performed to check statistical significance in growth of bacteria among different genotypes and indicated by alphabets e.g. a, b, c, d, e etc. which depict statistical difference from each other at *p-value* <0.001.

## Acknowledgements

IKD, MK and SB deeply acknowledge DST-SERB (Grant No: EMR/2016/001899) and Regional Centre for Biotechnology (RCB), Faridabad for providing financial support and central instrumental facilities. We express our gratitude to Prof. Walter Gassmann, Missouri University, USA, and Prof. Ashis Kumar Nandi, Jawaharlal Nehru University (JNU), New Delhi, India for providing *PstDC3000* strains and *sid2-1* lines and SA measurements, respectively used in this study. We also thank Dr. Souvik Bhattacharjee (JNU) for providing *p*ASK-IBA16 plasmid and Prof. Katsunori Tanaka, (Kwansei Gakuin University, Japan) for providing *E. coli* SUMOylation reconstitution system constructs. IKD acknowledges Department of Biotechnology (DBT), Government of India for providing fellowships during his PhD tenure and KIIT University, Bhubaneswar, India for his PhD registration. MK thanks UGC for his PhD fellowship.

## Author Contributions

SB conceived the research. IKD, MK and SB designed the research. IKD and MK performed all the experiments. HvdB generated *amiR-SUM2 sum3-1* line used in this study. SB helped in plasmid constructions and supervised the experiments. IKD, MK and SB analyzed the data and wrote the manuscript.

## Supporting information

Additional supporting information may be found in the online version of this article.

**S1 Fig: *sum3-1* partially alleviates, whereas *eds1-2* and *sid2-1* abolishes growth deficiencies of *sum1-1*.**

**S2 Fig: *sum1-1* and *sum2-1* plants are enhanced resistant whereas *sum3-1* is hypersusceptible to TNL-specific *PstDC3000(hopA1)* but not to CNL-specific *PstDC3000(avrRpm1)* avirulent strains.**

**S3 Fig: *sum1-1* and *sum2-1* leaves display increased whereas *sum3-1* is deficient in callose deposition in response to virulent *PstDC3000(EV)* or avirulent *PstDC3000(avrRps4)* infections.**

**S4 Fig: Developmental phenotype of 4-week-old SD grown plants of indicated genotypes.**

**S5 Fig: *sum1-1* and *sum2-1* display rapid whereas *sum3-1* is delayed than Col-0 in induction of *PR1* and *PR2* transcripts in response to virulent *PstDC3000(EV)* or avirulent *PstDC3000(avrRps4)* infections.**

**S6 Fig: Increased trichome density, elevated expression of PTI markers *FRK1* or *WRKY29*, and enhanced resistance to avirulent *PstDC3000(hopA1)* in *sum1-1* is SA-modulated.**

**S7 Fig: Enhanced basal immunity and ETI defences in *sum2-1* to TNL-specific *PstDC3000* strains is EDS1-dependent.**

**S8 Fig: Developmental defects and enhanced TNL-specific immunity in *sum1-1* is partially *SUM3* regulated.**

**S9 Fig: *SUM3* buffers increased induction of *PR1* in *sum1-1* in response to *PstDC3000* challenges.**

**S10 Fig: Schematic representation of key amino acid conservation and divergences among three *Arabidopsis* SUMO isoforms suggest their functional overlaps/distinctions.**

**S11 Fig: Complemented *sum1-1* or *sum3-1* lines have wild-type levels of SUMO1/2 conjugates upon *PstDC3000* (EV) infection.**

**S12 Fig: SUMO1 and SUMO3 cause reciprocal enhancements of SUMO-conjugates in *E. coli* SUMOylation-reconstitution system.**

**S13 Fig: *SUM3* mutation reduces SUMO1/2-conjugate enhancements during heat-shock.**

**S1 Table: List of primers used in this study.**

