## Supplementary Information for "Contrasting functions of *Arabidopsis* SUMO1/2 isoforms with SUMO3 intersect to modulate innate immunity and global SUMOylome responses"

**Supporting information**

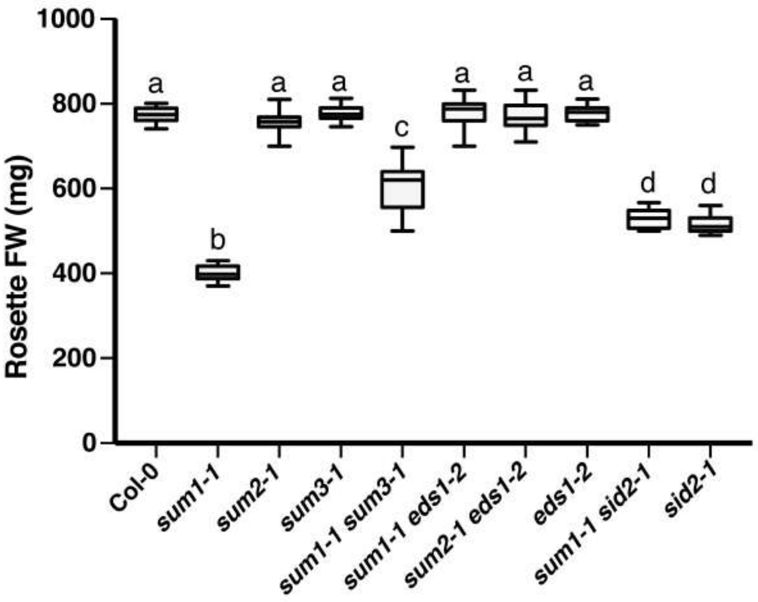


**S1 Fig: *sum3-1* partially alleviates, whereas *eds1-2* and *sid2-1* abolishes growth deficiencies of *sum1-1*.**

Fresh rosette weights (in mg) of 5-week-old SD grown plants of indicated genotypes were plotted into Whisker boxplot with Tukey test (n=20). ANOVA analysis was performed to identify statistical difference in rosette weights. The alphabets a, b, c and d define statistical significance (ANOVA) at *p-value* <0.001.

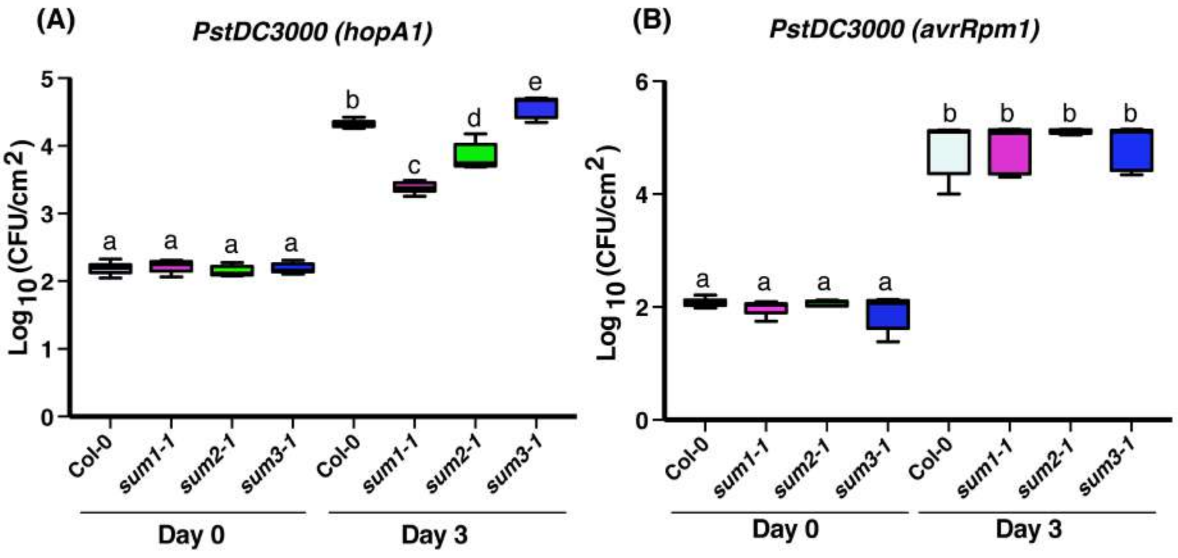


**S2 Fig: *sum1-1* and *sum2-1* plants are enhanced resistant whereas *sum3-1* is hypersusceptible to TNL-specific *PstDC3000 (hopA1)* but not to CNL-specific *PstDC3000 (avrRpm1)* avirulent strains.**

Bacterial infiltration of SD grown 4-week-old Col-0, *sum1-1*, *sum2-1*, and *sum3-1* plants were performed at a density of 5 X $10^4$ cfu $ml^{-1}$. Growth of bacteria was determined at indicated days and plotted into Whiskers box plot. Significance determined by ANOVA test with *p-value* <0.001 is indicated (n=6-9).

(A) Bacterial accumulation of *PstDC3000 (hopA1)* in indicated genotypes at Day 0 and Day 3 post-inoculation (dpi).

(B) Bacterial accumulation of *PstDC3000 (avrRpm1)* in indicated genotypes at Day 0 and Day 3 post-inoculation (dpi).

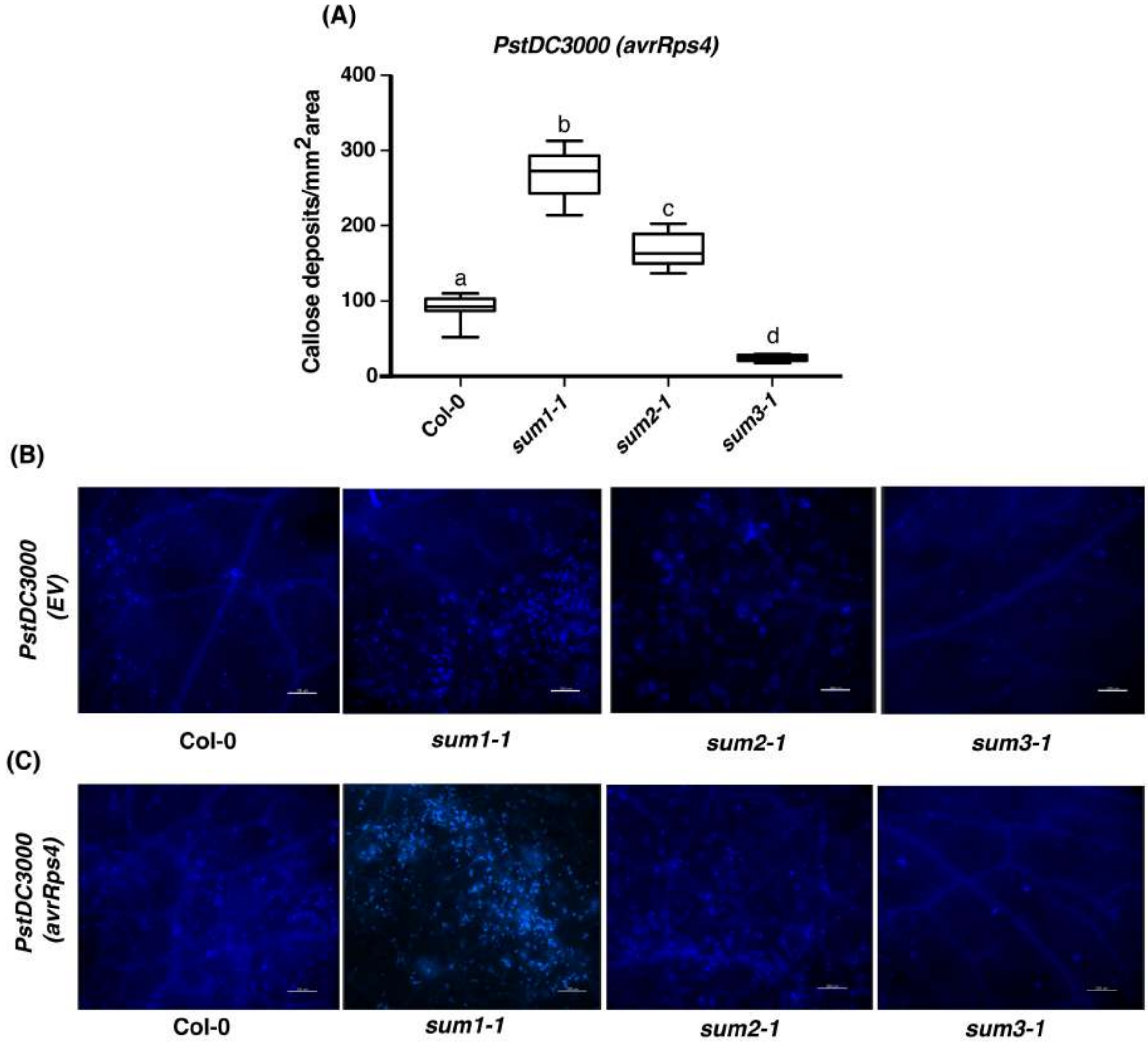


**S3 Fig: *sum1-1* and *sum2-1* leaves display increased whereas *sum3-1* is deficient in callose deposition in response to virulent *PstDC3000(EV)* or avirulent *PstDC3000(avrRps4)* infections.**

(A) Whisker boxplot of relative callose deposit densities on leaves of indicated plants infected with *PstDC3000* (*avrRps4*). For quantitation, callose deposits from 10 independent images (n=10) of same area were counted and plotted in a boxplot. The error bars indicate ANOVA at *p-value* <0.001.

(B) Representative images of callose deposits on leaves of indicated plants upon virulent *PstDC3000(EV)* infection.

(C) Representative images of callose deposits on leaves of indicated plants upon avirulent *PstDC3000(avrRps4)* infection.

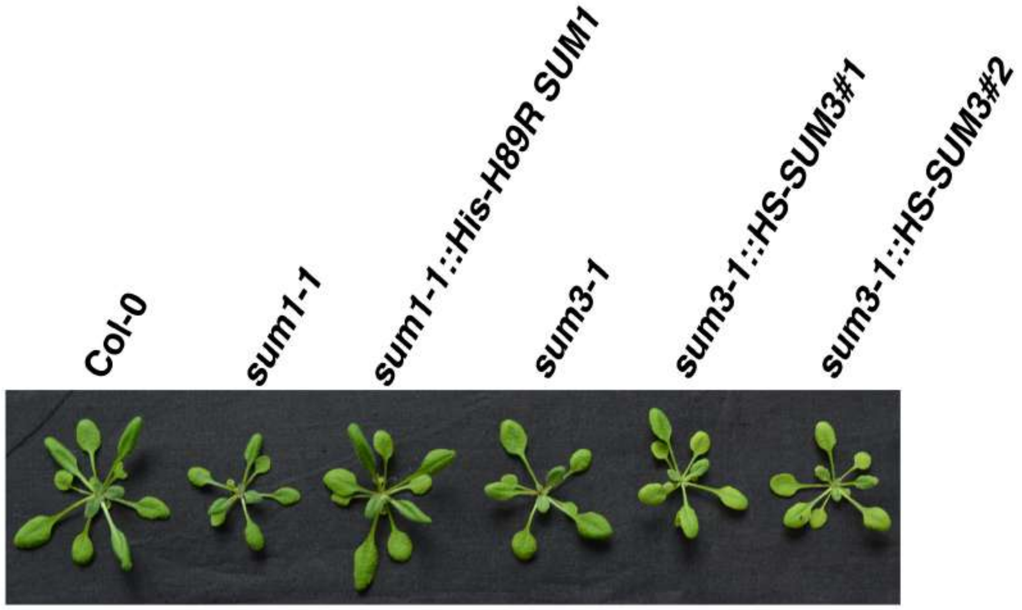


**S4 Fig: Developmental phenotype of 4-week-old SD grown plants of indicated genotypes.**

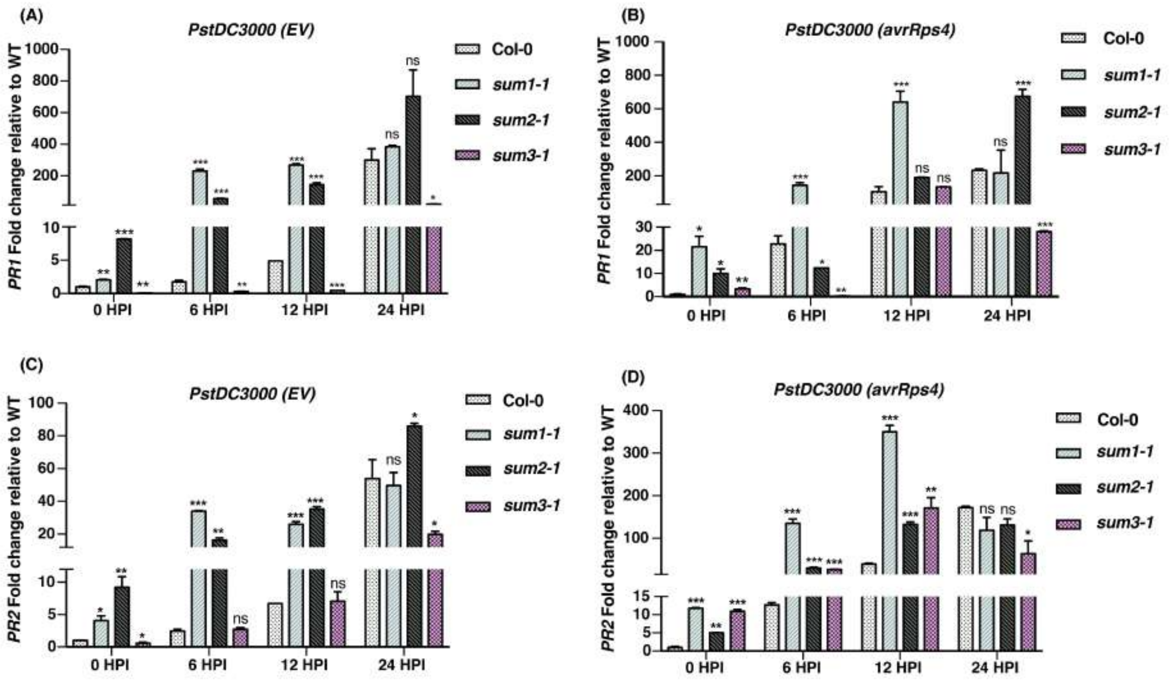


**S5 Fig: *sum1-1* and *sum2-1* display rapid whereas *sum3-1* is delayed than Col-0 in induction of *PR1* and *PR2* transcripts in response to virulent *PstDC3000(EV)* or avirulent *PstDC3000(avrRps4)* infections.**

*PstDC3000* infections, qRT-PCR analysis and data normalization were carried out as described in materials and methods. The Student's t-test was performed to calculate statistical significance with respect to expression levels in Col-0 at that particular time point (*=$p$<0.05; **=$p$<0.01; ***=$p$<0.001).

(A) Induction kinetics of *PR1* in response to virulent *PstDC3000 (EV)*.

(B) Induction kinetics of *PR1* in response to avirulent *PstDC3000 (avrRps4)*.

(C) Induction kinetics of *PR2* in response to virulent *PstDC3000 (EV)*.

(D) Induction kinetics of *PR2* in response to avirulent *PstDC3000 (avrRps4)*.

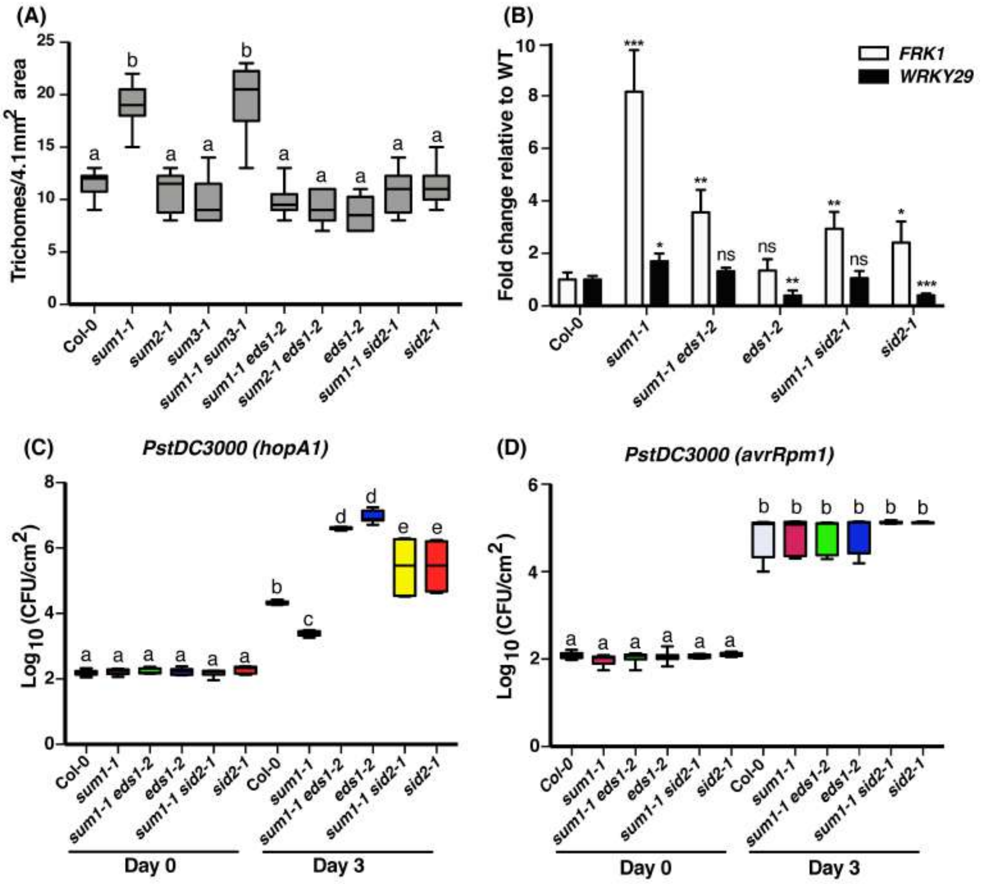


**S6 Fig: Increased trichome density, elevated expression of PTI markers *FRK1* or *WRKY29*, and enhanced resistance to avirulent *PstDC3000 (hopA1)* in *sum1-1* is SA-modulated.**

(A) Whisker Box plot of trichome densities on leaves of 4-week-old plants. Number of trichomes/ 4.1 $mm^2$ area were counted from images of leaf sections (n=10) observed under bright field in a fluorescence microscope. ANOVA was performed to check statistical significance among different genotypes at $p<0.001$ level.

(B) Fold change in basal expression levels of PTI markers *FRK1* and *WRKY29* in *sum1-1* and its combinatorial mutants. Expression levels of *FRK1* and *WRKY29* were determined by qRT-PCR, data normalization performed as described in materials and methods. The data is representative of three independent experiments. Error bars indicate SD (n=3). Student's t-test was performed to check statistical significance; *=$p$<0.05; **=$p$<0.01; ***=$p$<0.001, ns= not significant.

(C-D) Bacterial accumulation of *PstDC3000* (*hopA1*) or *PstDC3000* (*avrRpm1*) in indicated genotypes at Day 0 and Day 3 post-inoculation (dpi). Growth of bacteria was determined at indicated days and plotted into Whiskers box plot. Significance determined by ANOVA test with *p-value* <0.001 is indicated (n=6-9).

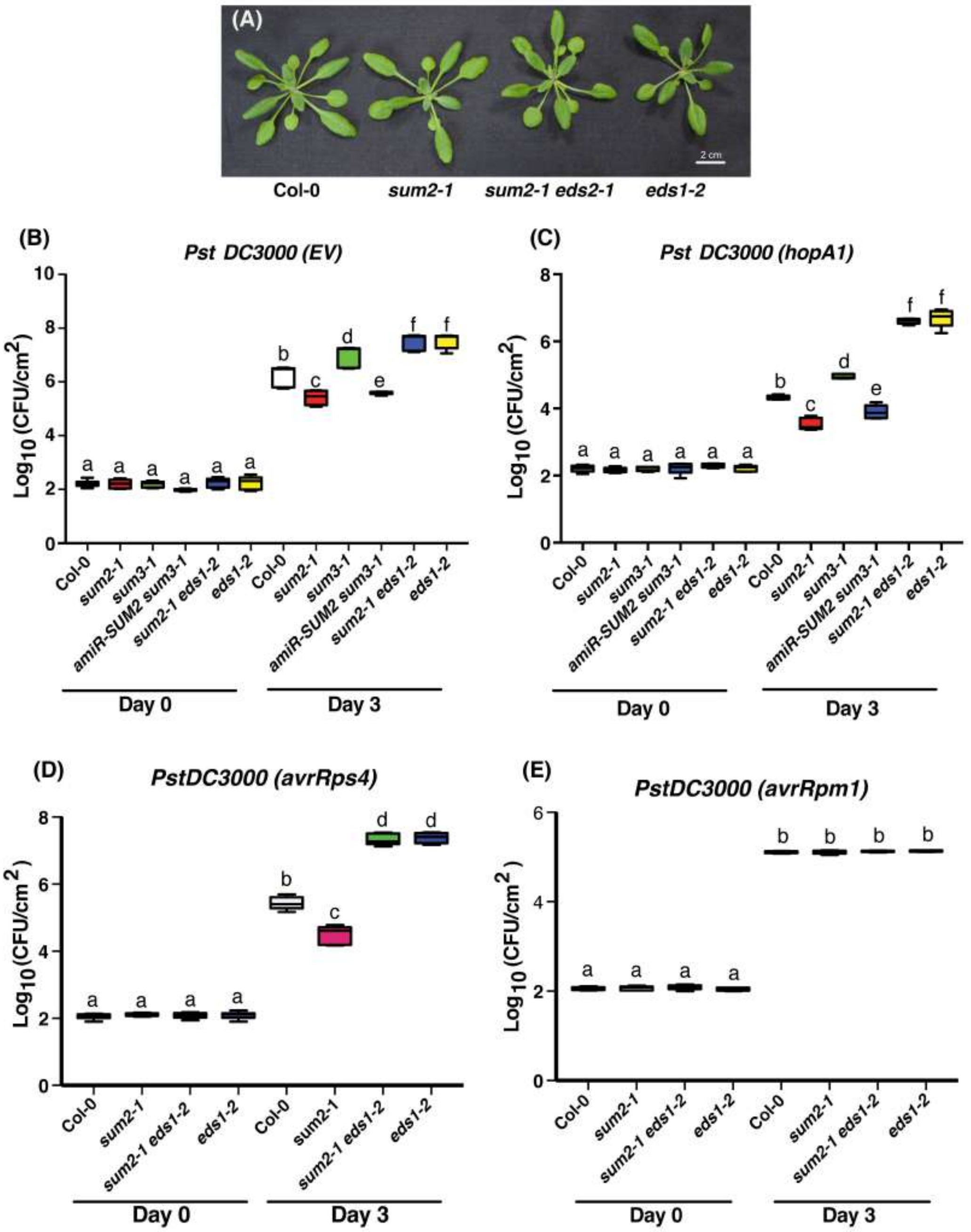


**S7 Fig: Enhanced basal immunity and ETI defences in *sum2-1* to TNL-specific *PstDC3000* strains is EDS1-dependent.**

(A) Developmental phenotypes of 4-week-old SD grown Col-0, *sum2-1, sum2-1 eds1-2* and *eds1-2* plants.

(B-E) Bacterial growth assays in Col-0, *sum2-1, sum3-1, amiR-SUM2 sum3-1, sum2-1 eds1-2* and *eds1-2* plants with *PstDC3000 EV* (B), *PstDC3000* expressing *hopA1* (C) *avrRps4* (D), and *avrRpm1* (E) effectors. Bacterial accumulation at Day 0 and Day 3 post-infiltration was determined as described in materials and methods. Values are plotted into whiskers box plot. Statistical differences determined by ANOVA test with *p-value* <0.001 is indicated (n= 6-9).

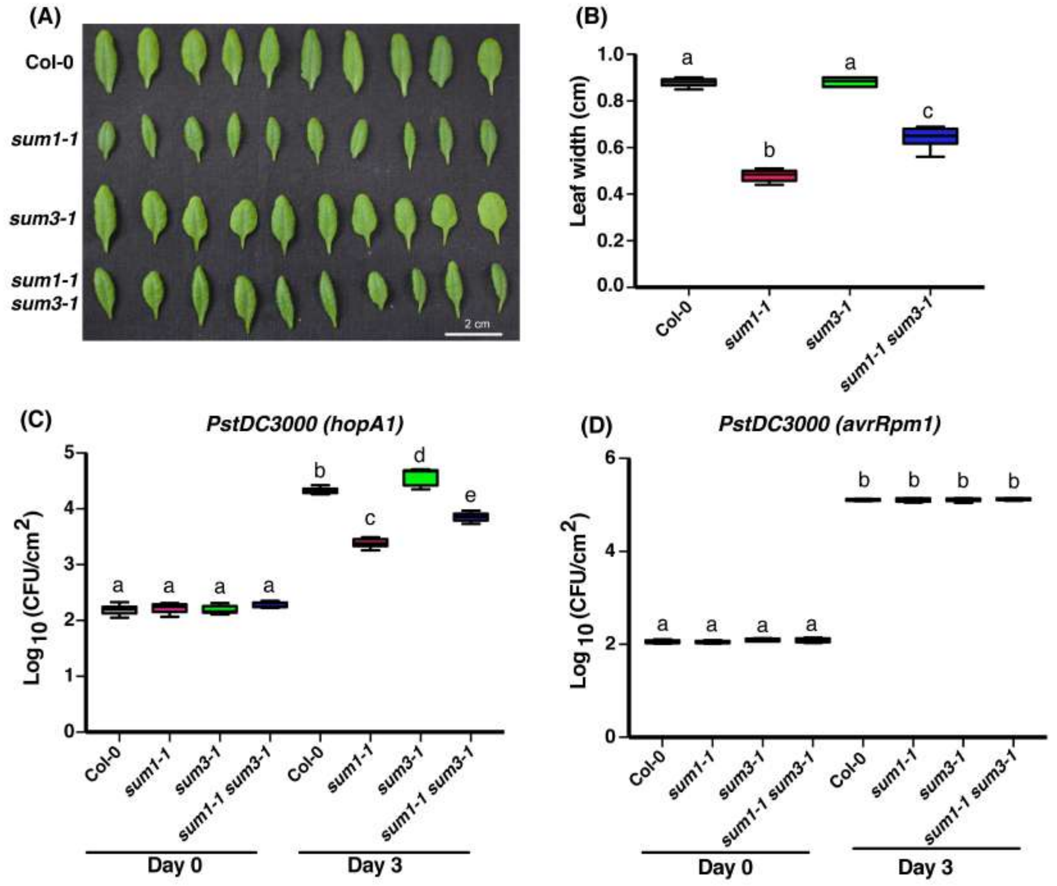


**S8 Fig: Developmental defects and enhanced TNL-specific immunity in *sum1-1* is partially *SUM3*-regulated.**

(A) Representative leaves from 4-week-old Col-0, *sum1-1, sum3-1* and *sum1-1 sum3*-1 plants grown under SD conditions.

(B) Whiskers boxplot of leaf widths (in cm) from indicated genotypes (n=10). Statistical differences are calculated at *p<0.001* level by ANOVA test.

(C-D) Bacterial growth assays on Col-0, *sum1-1, sum3-1* and *sum1-1 sum3-1* plants infected with avirulent *PstDC3000* expressing *hopA1* or *avrRpm1* effectors were performed as described earlier. ANOVA test was performed to calculate statistical differences at *p<0.001* level.

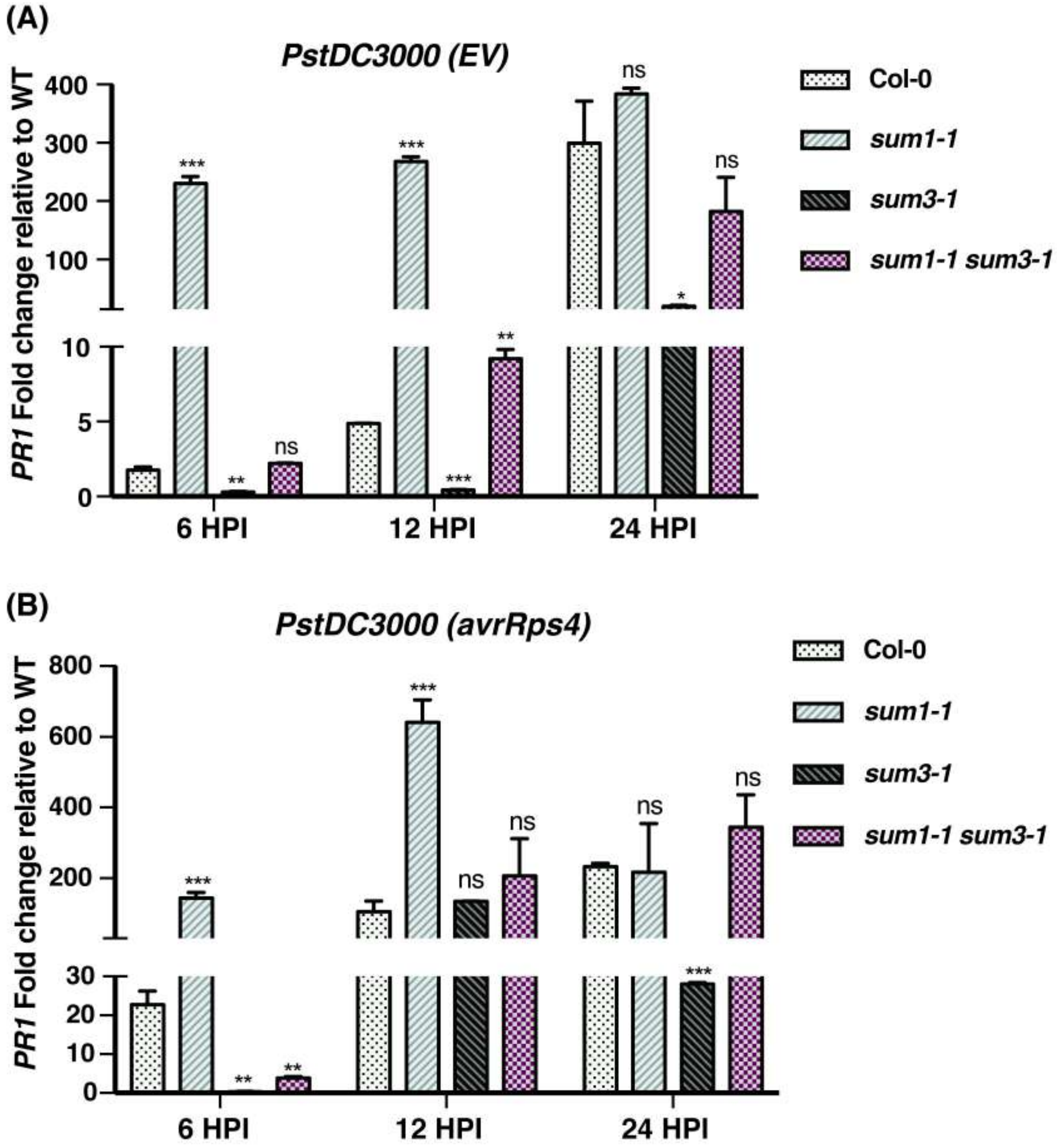


**S9 Fig: *SUM3* buffers increased induction of *PR1* in *sum1-1* in response to *PstDC3000* challenges.**

Leaves from 4-week-old SD grown plants were syringe infiltrated with indicated *PstDC3000* strains as described in materials and methods. *PR1* expression levels at 6, 12 and 24 hpi (hours post-infiltration) were determined. Student's t-test was performed to calculate statistical significance with respect to expression levels in Col-0 at that particular time point (*=$p$<0.05; **=$p$<0.01; ***=$p$<0.001, ns= non-significant).

(A) Induction kinetics of *PR1* in response to virulent *PstDC3000(EV)*.

(B) Induction kinetics of *PR1* in response to avirulent *PstDC3000(avrRps4)*.

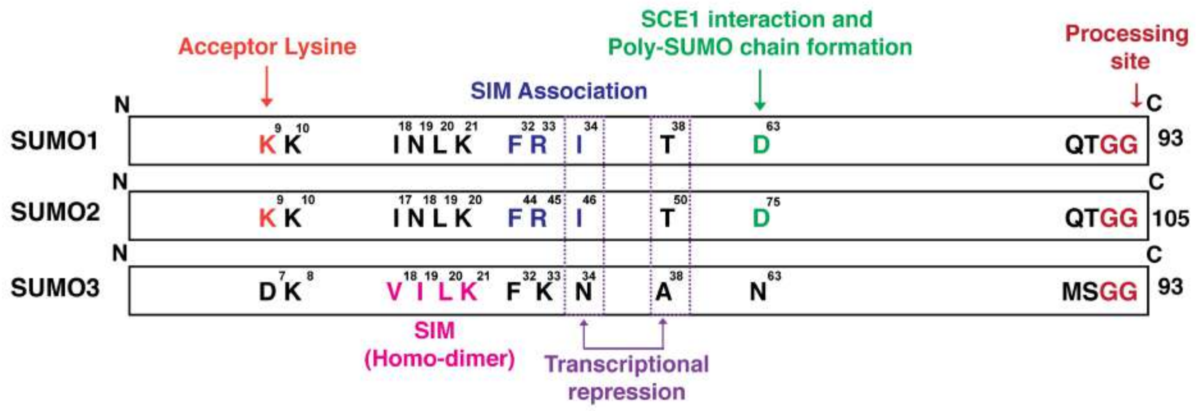


**S10 Fig: Schematic representation of key amino acid conservation and divergences among three *Arabidopsis* SUMO paralogs suggest their functional overlaps/distinctions.** SUMO1, SUMO2 and SUMO3 proteins are represented as rectangular boxes with selective amino acids and their implicated functional relevance are indicated.

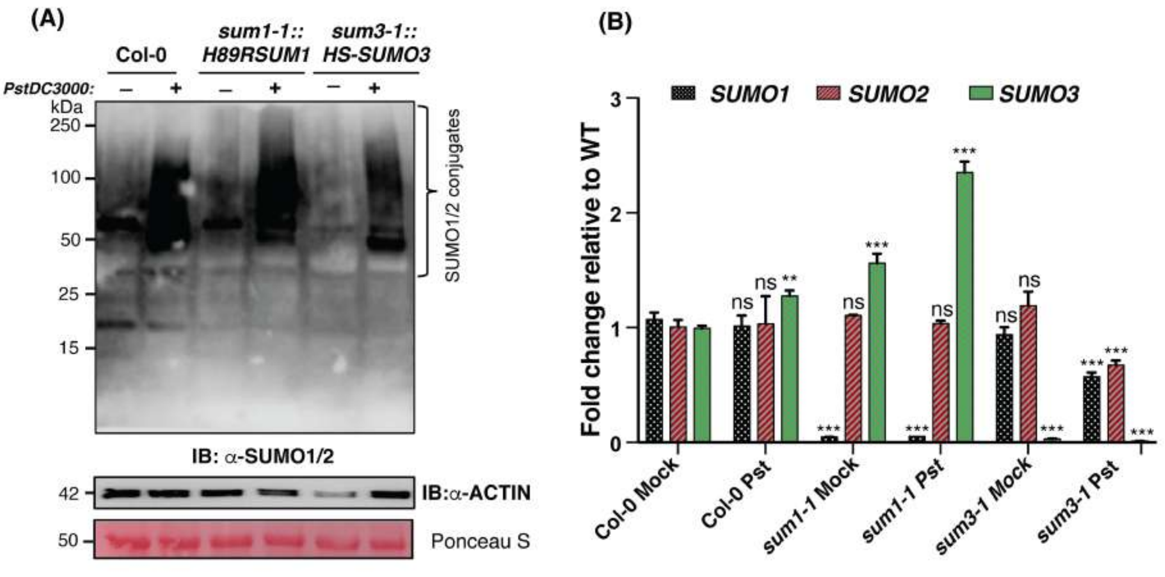


**S11 Fig: Complemented *sum1-1* or *sum3-1* lines have wild-type levels of SUMO1/2 conjugates upon *PstDC3000* (EV) infection.**

(A) The *sum1-1::H89RSUM1* and *sum3-1::HS-SUMO3#1* showed WT comparable levels of SUMO1/2 conjugates upon *PstDC3000 (EV)* infection at 24 hpi.

Total proteins extracts from 2-3-week-old seedlings subjected to **(A)** mock (10 mM $MgCl_2$) or *PstDC3000* (EV) infection were immunoblotted with anti-AtSUMO1 antibodies. Ponceau S stain of membrane for Rubisco subunit and immunoblot of ACTIN levels indicative of comparable protein loadings are displayed. Positions of molecular weight standards (in kDa) are indicated.

(B) Expression levels of *SUM* genes (fold change) were determined upon Mock (10 mM MgCl2) vs *PstDC3000 (EV)* infection at 24-hpi as described in materials and methods. The data is representative of three independent experiments. Error bars indicate SD (n=3). Student's t-test was performed to check statistical significance; *=$p$<0.05; **=$p$<0.01; ***=$p$<0.001, ns= not significant.

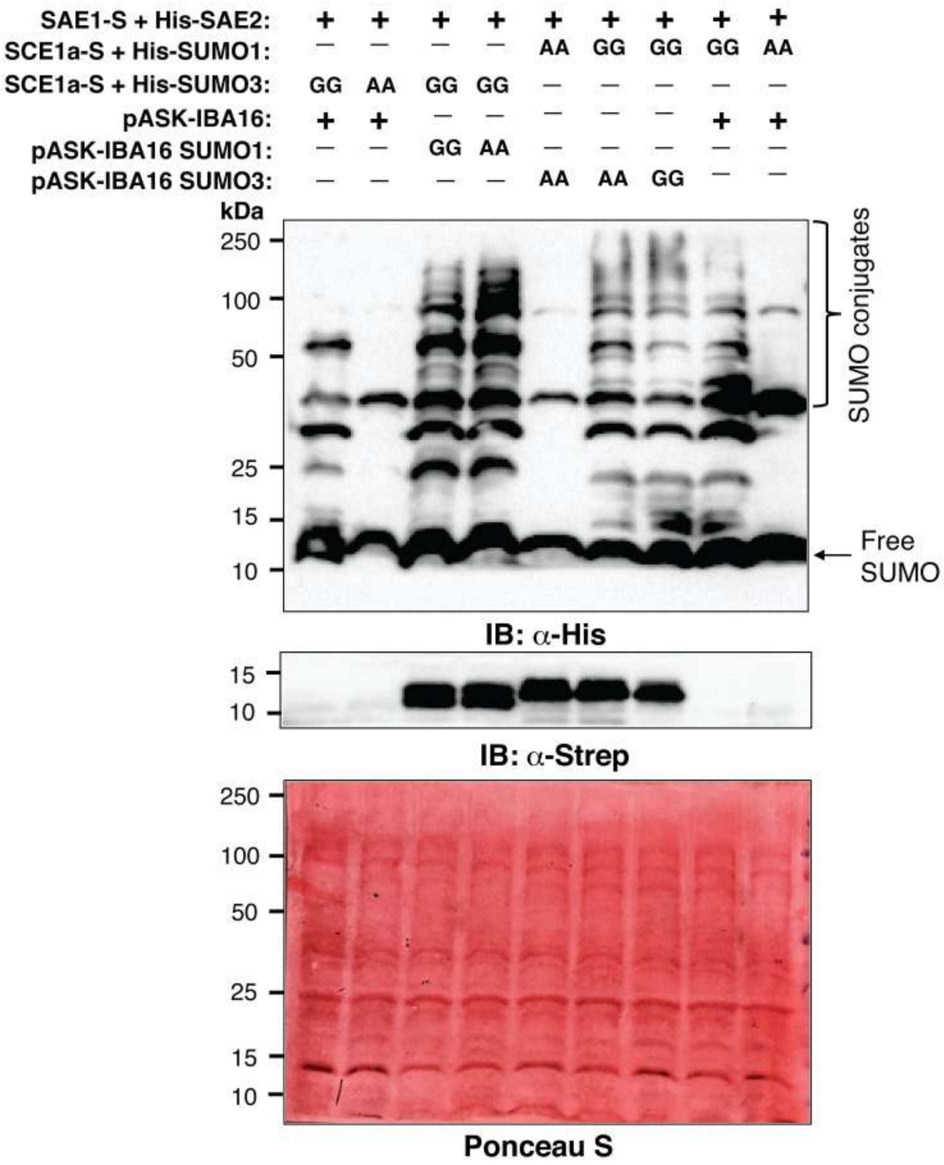


**S12 Fig: SUMO1 and SUMO3 cause reciprocal enhancements of SUMO-conjugates in *E. coli* SUMOylation-reconstitution system.**

*E. coli* BL21 (DE3) cells expressing the indicated combination of plasmids were induced as described in supplementary materials and methods. Cell lysates were immunoblotted with anti-His or anti-Strep antibodies as shown. The relative positions of SUMO-conjugates are depicted. The Ponceau S stained membrane is also shown to ensure equal protein loadings. Positions of protein molecular weight standards (in kDa) are indicated.

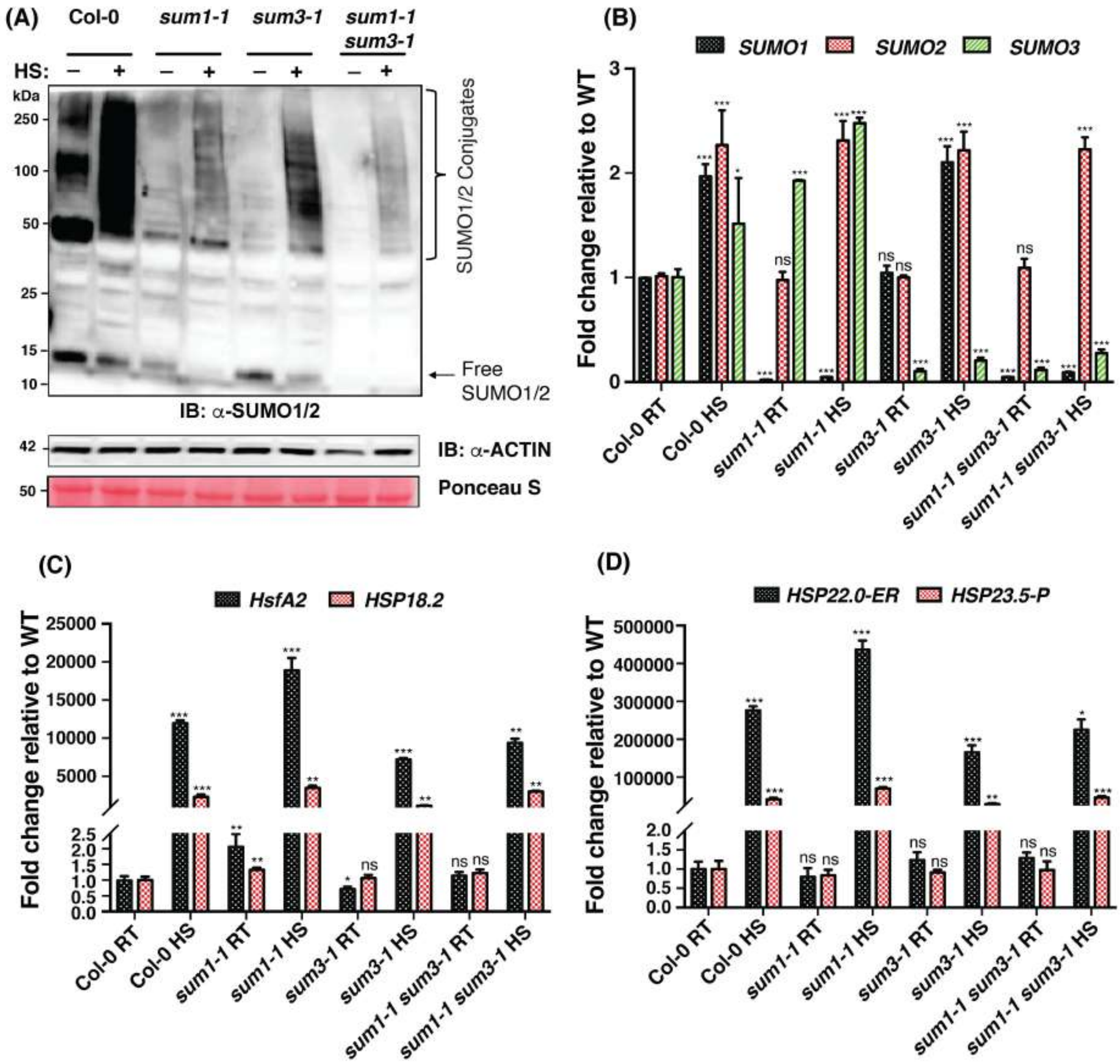


**S13 Fig: *SUM3* mutation alleviate the SUMO1/2-conjugate level upon heat shock.**

(A) Total proteins extracts from 15-days old seedlings subjected to $22^0$C (RT) or $37^0$C (HS; Heat shock) were immunoblotted with anti-AtSUMO1 antibodies. Ponceau S stain of membrane for Rubisco subunit and immunoblot with anti-ACTIN antibodies are indicative of comparable protein loadings. Positions of molecular weight standards (in kDa) are indicated.

(B-D) Expression levels of *SUMO* genes (B) and heat shock responsive genes-*HsfA2, HSP18.2* (C) and *HSP22.0-ER* and *HSP23.5-P* (D) in indicated genotypes were determined upon RT vs HS treatment as described in materials and methods. Error bars indicate SD (n=3). Student's t-test was performed to check statistical significance; *=$p$<0.05; **=$p$<0.01; ***=$p$<0.001, ns= not significant.

**S1 Table**

**Title: List of primers used in this study**

| Sl No | Primer Name | Primer sequence (5' to 3') | Used for |
|---|---|---|---|
| 1 | SUM1 Gateway For | AAAAAGCAGGCTCAATGTCTGCAA | Gateway cloning of SUM1GG/AA |
| 2 | SUM1GG Gateway Rev | AGAAAGCTGGGTATCAGCCACCAGTCTGATG | Gateway cloning of SUM1GG |
| 3 | SUM1AA Gateway Rev | AGAAAGCTGGGTATCAGGCAGCAGTCTGATG | Gateway cloning of SUM1AA |
| 4 | SUM3 Gateway For | AAAAAGCAGGCTCAATGTCTAACCCTCAAGATGA | Gateway cloning of SUM3GG/AA |
| 5 | SUM3GG Gateway Rev | AGAAAGCTGGGTAACCACCACTCATCGCCCGG | Gateway cloning of SUM3GG |
| 6 | SUM3AA Gateway Rev | AGAAAGCTGGGTATGCCGCACTCATCGCCCGG | Gateway cloning of SUM3AA |
| 7 | SAND qRT For | AACTCTATGCAGCATTTGATCCACT | To check gene expression |
| 8 | SAND qRT Rev | TGATTGCATATCTTTATCGCCATC | To check gene expression |
| 9 | PR1 qRT For | GTGGGTTAGCGAGAAGGCTA | To check gene expression |
| 10 | PR1 qRT Rev | ACCTTGGCACATCCGAGTCT | To check gene expression |
| 11 | PR2 qRT For | TCAAGGAAGGTTCAGGGATG | To check gene expression |
| 12 | PR2 qRT Rev | TTCACGAGCAAGGGAGATTG | To check gene expression |
| 13 | ICS1 qRT For | TACTAACCAGTCCGAAAGACG | To check gene expression |
| 14 | ICS1 qRT Rev | GAGGCTTGACAACAACTCTGT | To check gene expression |
| 15 | SUM1 qRT For | CTTGTTTGATGGGCGTCGTC | To check gene expression |
| 16 | SUM1 qRT Rev | CAGTCTGATGGAGCATCGCA | To check gene expression |
| 17 | SUM2 qRT For | TCGATGCAATGCTTCATCAGAC | To check gene expression |

| | | | |
|---|---|---|---|
| 18 | SUM2 qRT Rev | ACCGCCACTAAAAGCAGAAGA | To check gene expression |
| 19 | SUM3 qRT For | CGAGCAAATCAGCGTCAGTG | To check gene expression |
| 20 | SUM3 qRT Rev | ACACACACGATAACCGACCA | To check gene expression |
| 21 | SNC1 qRT For | TTGGAAGTCTCGAAGGGATG | To check gene expression |
| 22 | SNC1 qRT Rev | GTGGCCTTTGAAAGATCTGG | To check gene expression |
| 23 | FRK1 qRT For | CGGTCAGATTTCAACAGTTGTC | To check gene expression |
| 24 | FRK1 qRT Rev | AATAGCAGGTTGGCCTGTAATC | To check gene expression |
| 25 | WRKY29 qRT For | CCCGGAGAAATTCACCATAA | To check gene expression |
| 26 | WRKY29 qRT Rev | ATCAGCGGATGGGATCATAG | To check gene expression |
| 27 | sum1-1 LP | TTTCGTGTAGCTGCGATTAGG | For PCR-based genotyping of *sum1-1* mutation |
| 28 | sum1-1 RP | TTATCTTTGCTCGCCATTAGC | For PCR-based genotyping of *sum1-1* mutation |
| 29 | sum2-1 LP | GTCGGAGAATCGGATTTCTTC | For PCR-based genotyping of *sum2-1* mutation |
| 30 | sum2-1 RP | TGAGGGTGTGTATTGGTGGAG | For PCR-based genotyping of *sum2-1* mutation |
| 31 | sum3-1 dspm1 | CTTATTTCAGTAAGAGTGTGGGGTTTTGG | For PCR-based genotyping of *sum3-1* mutation |
| 32 | EDS1 F2 | CCCTTTCTAGTTTCCTTGAGCTAAG | For PCR-based genotyping of *EDS1*/*eds1-2* |
| 33 | EDS1 R3 | TCAGGTATCTGTTATTTCATCCATC | For PCR-based genotyping of *EDS1*/*eds1-2* |
| 34 | LB3 | TAGCATCTGAATTTCATAACCAATCTCGATACAC | For PCR-based genotyping of mutants from SAIL stock |
| 35 | SALK_LB1.3 | ATTTTGCCGATTTCGGAAC | For PCR-based genotyping of mutants from SALK stock |
| 36 | Sm108F | TTCTTCATGCAGGGGAGGAG | For PCR-based genotyping of *SID2*/*sid2-1* |
| 37 | Sm30F | CAACCACCTGGTGCACCAGC | For PCR-based genotyping of SID2/*sid2-1* |
| 38 | L1849R | AAGCAAAATGTTTGAGTCAGCA | For PCR-based genotyping of SID2/*sid2-1* |

| | | | |
|---|---|---|---|
| 39 | KpnI-SUM3p For | GGGGTACCAAAATTAACAAAGGTACAAAGATTCT | For PCR-amplification of *SUM3* genomic sequence |
| 40 | XbaI-SUM3-UTR Rev | GGTCTAGAAATAACCATGCATGCATTAGAAGTTTA | For PCR-amplification of *SUM3* genomic sequence |
| 41 | His-StrepII Rev | AGGATGAGACCATGAACCACCTCCTCCACCGTGA TGGTGATGGTGATGCATCTTTCCTTTTATCAGAT | For insertion of His-StrepII Tag in *SUM3* genomic sequence |
| 42 | His-StrepII For | AAGATGCATCACCATCACCATCACGGTGGAGGAG GTGGTTCATGGTCTCATCCTCAATTTGAAAAATG ATGTCTAACCCTCAAGAT | For insertion of His-StrepII Tag in *SUM3* genomic sequence |
| 43 | HsfA2 qRT For | TGGGATTCTCATAAGTTCTCAACA | To check gene expression |
| 44 | HsfA2 qRT Rev | TGGATCAATCTTTCTGAATCCAT | To check gene expression |
| 45 | HSP18.2 qRT For | TTACCGAATGCAAAGATG | To check gene expression |
| 46 | HSP18.2 qRT Rev | CGGAGATATCGATGGACTTGA | To check gene expression |
| 47 | HSP22.0-ER qRT For | ACTACTCCAGGCAGCTTGCTA | To check gene expression |
| 48 | HSP22.0-ER qRT Rev | CTTGAATGGATCAGGGAACC | To check gene expression |
| 49 | HSP23.5 qRT For | GATCAAGATGCGTTTCGACAT | To check gene expression |
| 50 | HSP23.5 qRT For | TTCTACAGAGATTTTGACGTCTTCTT | To check gene expression |
| 51 | SAE1 qRT For | ATTCCTCGGAGAACAGCAAA | To check gene expression |
| 52 | SAE1 qRT Rev | ACGCCCTTCACTCTCTTCAA | To check gene expression |
| 53 | SAE2 qRT For | ACAGCCTTTTTGAAGCGAAA | To check gene expression |
| 54 | SAE2 qRT Rev | ATAATGGCGTTGGTCGTAGC | To check gene expression |
| 55 | SCE qRT For | AATGGTGTGGCATTGCACTA | To check gene expression |
| 56 | SCE qRT Rev | TCCTCACTGAAGTGCATCGT | To check gene expression |

| | | | |
|---|---|---|---|
| 57 | SIZ1 qRT For | AACAGGGAAAGAAGCAGGAA | To check gene expression |
| 58 | SIZ1 qRT Rev | GGCAGCTTGTTCATCAGAAA | To check gene expression |
| 59 | HPY2 qRT For | CTACACCTTCCTCAGTGCCA | To check gene expression |
| 60 | HPY2 qRT Rev | AACATTCCAAACTGCTTCCC | To check gene expression |
| 61 | ESD4 qRT For | TGATGCTGGATTCGTTGTCG | To check gene expression |
| 62 | ESD4 qRT Rev | TCTCCGCACTTTGCATAAGC | To check gene expression |
| 63 | ELS1 qRT For | TGGATGGTTACCACAAACGG | To check gene expression |
| 64 | ELS1 qRT Rev | ATGAACTTGTCTCCGCAACG | To check gene expression |
| 65 | OTS1 qRT For | TCACGGTCAACATTTTCTGC | To check gene expression |
| 88 | OTS1 qRT Rev | CAGGCTTTCTACCAGCCTTG | To check gene expression |
| 67 | OTS2 qRT For | GCCTCAAAAGACACCTCGG | To check gene expression |
| 68 | OTS2 qRT Rev | GCTTATCCAGCTTCCACGTC | To check gene expression |
